# A hierarchical Bayesian model for dynamic phenotypes in multi-environment trials with genotype by environment interaction

**DOI:** 10.64898/2026.11.05.764527

**Authors:** Shota Gugushvili, George A. K. van Voorn, Martin P. Boer, Daniela Bustos-Korts, Sandra Huynh Truong, Nicholas A. Friedenberg, Ryan McCormick, Scott Chapman, Carlos D. Messina, Fred A. van Eeuwijk

**Affiliations:** Mathematical and Statistical Methods group – Biometris, Wageningen University and Research, Wageningen, The Netherlands; Institute of Plant Production and Protection, Faculty of Agricultural Sciences, Universidad Austral de Chile, Valdivia, Chile; Research & Development, Corteva Agriscience, Johnston, IA, United States; School of Agriculture and Food Sustainability, Faculty of Science, The University of Queensland, Gatton, QLD, Australia; Department of Horticultural Sciences, University of Florida, Gainesville, FL, United States

**Keywords:** crop growth model, genotype by environment interaction, phenomics, phenotyping, process-based model, yield prediction

## Abstract

New phenotyping tools produce longitudinal phenotypes that describe plant growth and development. Estimates for the parameters that describe the dynamics of such longitudinal phenotypes can be incorporated as predictors in models for static target traits like yield and biomass. Within a hierarchical Bayesian context, we present a differential equation (DE) model for log-logistic growth in longitudinal traits for sets of genotypes across multiple environments. The DE parameters coincide with a midpoint, a slope, and an asymptote at the end of growth and development, i.e., final biomass. Each DE parameter in our hierarchical model results from contributions of genotype, environment, and *G* × *E*. This formulation allows one to assess and quantify the genotype-specificity of the DE parameters. When none of the DE parameters in our model requires a *G* × *E* term, the dynamic parameters characterizing the particular trait will be useful features for inclusion in prediction models for yield or biomass. We fitted a DE model to APSIM-simulated longitudinal wheat biomass data and quantified the contribution of genotype, environment and *G* × *E* to the phenotypic variation in the DE parameters. Only the upper asymptote showed clear *G* × *E*. Estimates for genotypic and *G* × *E* parameters within the DE parameters were used to characterize genotypes. Estimates for environmental and *G* × *E* effects formed inputs for environmental characterization.

## 1 Introduction

New phenotyping devices produce longitudinal data on many aspects of plant growth and development for large collections of genotypes (Guédon et al., 2022). Depending on the device, these data have moderate to high time resolution (Araus et al., 2018, 2022; Tardieu et al., 2017). The study of longitudinal phenotypic traits is of interest in itself when these traits can be considered to represent yield components, i.e. they bear a physiological relation to yield and reflect adaptation of the genotype to the environment. Knowledge of the dynamics of yield components adds to biological insights in yield-producing processes and will improve yield prediction models. Such improvements can cover both within-trial yield forecasts as well as between-trial yield predictions where the longitudinal phenotyping information can be combined with genomic information.

Beyond a small subset of longitudinal traits that can be considered to be directly or closely related to yield, most longitudinal traits that are measured automatically or semi-automatically by proximate and remote sensors initially lack a direct connection with plant physiological processes and structures. Such traits are called secondary traits, and although they lack clear biological meaning, it can be worthwhile to model their dynamics and estimate the corresponding parameters because aspects of their dynamical behavior may improve prediction models for yield. When secondary traits or functions of secondary traits contribute to the prediction of yield, then it is plausible that they represent proxies of physiologically important traits, that is, yield components. Various articles offer guidelines on how to decompose yield into components and which traits to phenotype as proxies to those components (Araus et al., 2018; Tardieu et al., 2017; Bouidghaghen et al., 2023). These yield components characteristically reflect resource capture and conversion into biomass and aspects of source-sink relations that lead to the prediction of the fraction of biomass translated into yield (harvest index). A decomposition of yield into component traits or processes immediately suggests yield prediction models that integrate information from multiple yield components as measured by proxies. The yield components act as genotype-specific inputs. The prediction model consists of a function that is able to convert the yield components together with some environmental inputs into yield as output. Prediction models differ with respect to the yield components to be included as static or dynamic quantities. The static yield components represent physiological constants that may be observed in single well-conditioned experiments, while the dynamic yield components require longitudinal observations. The environmental inputs are dynamical, but may differ at the aggregation level that is incorporated. For example, temperatures may be chosen to be included as a daily or weekly mean.

Bouidghaghen et al. (2023) make a case for the breakdown of yield into a number of subprocesses and structures, each of which can be measured by high-throughput robotized phenotyping devices at platforms. They emphasize that before inserting traits measured on platforms into prediction models for yield in the field, these traits require scaling, transformation, and conditioning on environmental variables such as, for example, thermal time, soil water potential, and vapor pressure deficit. Although Bouidghaghen et al. (2023) illustrate how to decompose yield and predict yield components from phenotyping and genomics, they do not go all the way in formulating and comparing prediction models for yield from genotype-specific and environmental inputs. The same approach to yield prediction via yield decomposition and component approximation drives the work of other researchers that aims at the integration of longitudinal phenotyping information and ultimately genomics into yield prediction models. Examples of this approach can be found in Roth et al. (2021, 2022, 2023) and Pérez-Valencia et al. (2022, 2024). All of these papers focus on one or a few longitudinal traits whose dynamics are modeled to extract summary statistics for eventual prediction of yield in future environments. Again, this latter aspect is not explicitly elaborated.

The papers mentioned in the above paragraphs all depart from dynamical models for yield (or yield components) that are based on physiological insights and help to define underlying yield components that can either be measured by sensors in the field or on a platform or predicted from marker profiles in an attempt to integrate phenotyping with genomics in the prediction of yield. The dynamical model can be a fully developed ecophysiological model, such as in Technow et al. (2015) and Messina et al. (2018), or more data driven as in the case of Roth et al. (2021, 2022) and Pérez-Valencia et al. (2022, 2024). In all cases, an important consideration in the use of longitudinal phenotypic information for yield prediction is that the dynamics of the longitudinal trait itself, say the defining parameters, are not subject to strong genotype by environment interaction, *G* × *E*, themselves (Poudel et al., 2023; Roth et al., 2023). The idea here is that parameters describing longitudinal yield components will enter as genotype-specific static inputs into yield generating functions together with dynamical environmental inputs. When genotype-specific inputs do not show *G* × *E*, or when they show a simple type of *G × E* that can be scaled, transformed, or conditioned on an environmental covariable (Bouidghaghen et al., 2023), they can be measured or estimated in a single platform or field experiment. *G* × *E* for yield then needs to emerge from the yield generating function.

The objective of this paper is to investigate a class of statistical models for analyzing longitudinal trait data for a population of genotypes observed by one or more phenotyping devices in a series of environments. The longitudinal traits may have been chosen after a prior yield decomposition and the expectation is that the parameters describing the longitudinal behavior of the component trait will exhibit relatively little *G* × *E* and that they will ultimately contribute to a better yield prediction. For improved yield prediction, we may later need to combine dynamic information on multiple yield components and environmental inputs, but on the road to a robust predictive model, we first need to develop adequate models for describing individual longitudinal traits. We aim for a modeling framework for our individual longitudinal traits that is flexible in the kind of functions and parameterizations that can be chosen to describe the dynamics of the trait. Simultaneously, across the entire set of genotypes and environments, we would like to test and quantify the amount of *G* × *E* in each of our dynamic parameters, using the same inferential scheme that was used to estimate these parameters.

We will elaborate here the modeling concepts for plant growth that were introduced in Van Voorn et al. (2023). In that paper, simple differential equation (DE) models with few state variables and parameters were presented that were derived from first principles. Subsequently, these models were fitted per genotype and per trial, that is, the estimation process was done separately for each combination of genotype and environment without borrowing information between genotypes or environments, although it can be assumed that a population of genotypes will show similar growth patterns over time just as environments with comparable stress will do. Fitting dynamic models for single genotype by environment combinations is feasible when the time resolution for observations is high, say, daily to a few times per week (Roth et al., 2021, 2022; Shao et al., 2026). However, such a resolution is not realistic for most field situations. Longitudinal trait data produced by drones in outside-field experiments typically consist of series of 10 to 15 time points in a growing season. For such medium-resolution temporal data, borrowing information between genotypes in the fitting process will be beneficial, and very likely the same is true for borrowing information between environments. With medium-resolution secondary phenotypes in mind, we present here a hierarchical Bayesian framework for the simultaneous fitting of dynamic models on longitudinal trait data for populations of genotypes across multiple environments. This framework allows borrowing of information between genotypes and environments and has as further advantages the use of shrinkage estimation of the dynamical parameters as well as appropriate propagation of uncertainty on parameter estimates and phenotypic predicted values. As the most distinguishing feature, our framework expresses dynamic parameters as functions of genotypic, environmental, and *G* × *E* terms in a fashion reminiscent of generalized linear mixed models (GLMMs). The decomposition of the phenotypic variation of the dynamic parameters into components attributable to genotype, environment, and *G* × *E* provides information on the genotype specificity of these parameters and the genotypic and environmental drivers of dynamic behavior. An example analysis on APSIM simulated wheat (*Triticum aestivum* L.) biomass data, the generation of which is described in Bustos-Korts et al. (2019a,b), illustrates the machinery of our Bayesian hierarchical approach together with the interpretation of the estimates of the genotypic and environmental contributions to the DE parameters.

## 2 Materials and methods

In this section, we present key aspects of our approach. We first give a rough and intuitive sketch of our approach and philosophy. We then recall a phenomenological model from Van Voorn et al. (2023) and show how a modification of it can be embedded within a statistical model. The resulting model will be fitted via a Bayesian approach. An APSIM simulated wheat dataset is used to illustrate and investigate the practical performance of our method.

### 2.1 A sketch of our hierarchical approach

In a generic notation, we can write the signal part of the dynamics of the *s*^*th*^ secondary trait for a genotype, *i*, in an environment, *j*, as:

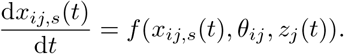

Here *x*_*ij*,*s*_(*t*) represents, for genotype *i* in environment *j*, an error free time-dynamic secondary trait, *s*, such as biomass. The variable *z*_*j*_(*t*) describes a vector of time-dynamical exogenous inputs, such as one or more soil properties, minimum, mean and maximum temperatures, and precipitation. The DE or process parameters are contained in the vector *θ*_*ij*_. From this ingredient we build a nonlinear mixed model, in which each DE parameter is related to a linear mixed predictor through a log link, in the spirit of a GLMM. In particular, the hierarchical system we propose can be thought of as containing an extra level of modeling in defining GLMMs for each of the DE parameters: *g*(*θ*_*ij*_) = *Xγ*, where the link function *g*(·) works on the expectation of *θ*_*ij*_ and is linear in the regression parameters *γ*, with design matrix *X*. The GLMMs for the DE parameters will include a general intercept, a genotypic main effect term, an environmental main effect term, and a *G* × *E* term. This formulation allows testing explicitly for the inclusion of *G* × *E* terms in the GLMMs for the individual DE parameters. Subsets of genotypes and environments for whose dynamics no *G* × *E* terms are needed act in what we can call a genetically additive way, i.e., it suffices to estimate the differences between genotypes in a single environment because the same genotypic differences will occur in any environment in which the genotypes will be grown, i.e. predictions are valid for any environment. To obtain a genomic prediction model, the genotypic main effects can be replaced or approximated by penalized regressions on marker profiles. It is important to note that additivity of genotype and environment for the DE parameters does not exclude *G* × *E* at the level of the dynamic trait itself because the non-linearity of the DE system can still create *G* × *E* as an emerging property on top of DE parameters that behave additively with respect to genotype and environment.

### 2.2 Phenomenological model

Van Voorn et al. (2023) discuss four models that reflect some essential characteristics of crop growth. These models are of low complexity, while being time-dynamical, as well as phenomenological in nature, and offer a biological interpretation of their parameters. According to Van Voorn et al. (2023), genotypic variation can be conceptualized as genotype-related differences in parameter values, while dependence on environmental or management inputs can be explicitly introduced by making the model parameters functions of such inputs.

In this paper, we focus on a logistic model for biomass in wheat. This is the most basic model in Van Voorn et al. (2023), which we will combine with a hierarchical Bayesian modeling approach. The resulting statistical model is flexible enough to reach non-trivial inferential conclusions. In addition, the logistic model serves as a useful baseline against which other approaches can be benchmarked and compared. Furthermore, there is room to explore common numerical challenges in nonlinear regression and fitting of ordinary differential equation (ODE) models (Seber and Wild, 1989; Ramsay and Hooker, 2017), as well as to examine the scalability of the method to large data.

#### 2.2.1 Logistic model

In the notation of Van Voorn et al. (2023), the logistic model is given by the ODE

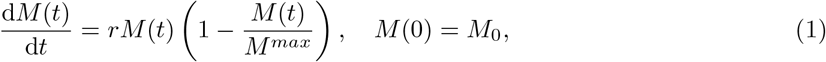

with an explicit solution

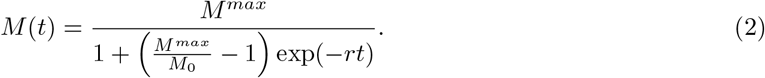

Here *M* (*t*) is the total biomass at time *t*, the positive parameter *r* is an intrinsic growth rate, while the positive parameter *M*^*max*^ is an intrinsic growth limitation or asymptote. Depending on the context, the initial condition *M*_0_ can be interpreted either as the seed weight, or as the biomass at the starting day of measurement.

In the literature there exist several alternative parameterizations of the logistic model (1)–(2), see, e.g., Seber and Wild (1989) and Pinheiro and Bates (2000). In one parameterization, the three parameters of the model are the intrinsic growth rate *r*, the intrinsic growth limitation or asymptote *M*^*max*^, and the midpoint.

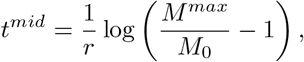

where the biomass *M* (*t*^*mid*^) = *M*^*max*^*/*2. Here we implicitly assume that *M*_0_ *< M*^*max*^. The function itself is then given by

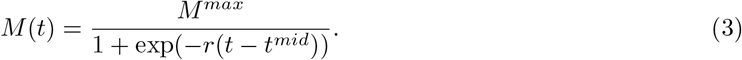

This parameterization will be the basis for our subsequent arguments.

#### 2.2.2 Log-logistic model

Successful fitting of nonlinear regression models to the data is strongly dependent on the chosen parameterization of the function to be fitted (Seber and Wild, 1989, pp. 673–674). For example, a reparameterization of a nonlinear model could remedy parameter identifiability (Seber and Wild, 1989, p. 103). A suitable reparameterization of a model could likewise improve the convergence of an iterative non-linear optimization algorithm (Seber and Wild, 1989, pp. 95–97), or, in the context of the present paper, convergence of an MCMC sampler. Ross (1990) contains further discussion on these topics.

A modified version of the logistic model (1)–(2), termed the log-logistic model, turned out to be numerically easier and faster to fit in the hierarchical modeling context than the basic logistic model. This appears to be caused by algorithmic peculiarities of our MCMC sampler. In general, both the logistic and log-logistic curves can be expected to give visually quite similar fits.

The log-logistic growth function closely resembles the logistic model (3) and is given by

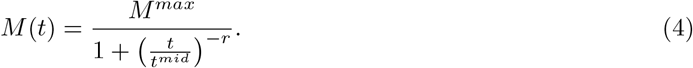

The three parameters of the log-logistic model have an intuitive interpretation similar to the parameters of the logistic model: *M*^*max*^ is the intrinsic growth limitation. Next, *t*^*mid*^ is the time point where response *M* (*t*^*mid*^) = *M*^*max*^*/*2. Finally, the parameter *r* is the scaled slope at the midpoint. See Ritz et al. (2020) for additional details.

The log-logistic model is by far the most common dose response model (Ritz et al., 2020, p. 178). Its name reflects the fact that the function is obtained by rescaling the log-logistic distribution, which in economics is known as the Fisk distribution. The log-logistic distribution also arises in the survival analysis context (Bennett, 1983). Its mathematical properties make it suitable for crop growth modeling as well, and in fact the model has been used in the agricultural setting: some references are listed in Focke et al. (2017).

The differential equation for the log-logistic growth model away from the time origin *t* = 0 can be shown to be

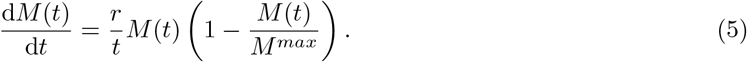

The initial condition is *M* (*t*^*mid*^) = *M*^*max*^*/*2, and the equation is thus understood to operate forward and backward in time. When fitting the log-logistic model to data, we will rely on the explicit form (4) rather than the differential equation formulation (5). See Section 2.5 ahead.

### 2.3 Data description

To illustrate our method, we chose the same simulated longitudinal wheat data as in Van Voorn et al. (2023). The crop growth simulation platform was APSIM-Wheat (Keating et al., 2003; Holzworth et al., 2018).The data consist of biweekly (i.e. successive measurements spaced two weeks apart) biomass observations over the growing season for an Australian diversity panel with originally 199 wheat genotypes (Zheng et al., 2013) observed at four locations over 31 growing seasons (years); see Bustos-Korts et al. (2019a) and Bustos-Korts et al. (2019b). The biweekly observations were chosen to reflect typical medium-resolution temporal data that are similar to drone imagery observations for biomass. The complete data set is available online (Bustos-Korts, 2023). For computational convenience, we drew a genetically representative subset of 48 genotypes from the entire set of 199 genotypes (see the code referred to in the Software section). This subsetting of genotypes allowed us to do all calculations described in this paper on a laptop (MacBook Air, 2020, with Apple M1 chip with 8 cores, 4 for performance and 4 for efficiency, and 8 GB RAM). The four locations chosen (Emerald, Narrabri, Yanco, Merredin) corresponded to environmentally contrasting wheat growing areas in the Australian wheat belt. The environments in this paper are a subset of the environments described in Casadebaig et al. (2016) and Chenu et al. (2013).

The first 14 days of biomass data after sowing were discarded, as these biomasses were (near) zero, which presented numerical problems for fitting. The biomass given in kg ha^*−*1^ was converted into tons ha^*−*1^, and the time in days was re-scaled by a factor of 10, so that a new time unit corresponded to 10 days. These transformations bring both variables to the same scale and can be expected to help with non-linear fitting, for instance because the population level parameters are then of the same order of magnitude (Seber and Wild, 1989, p. 674).

At this stage of research, we preferred simulated data over field trial data. This choice allows for an objective assessment of our approach against the ground truth. On the environmental side, the data were qualitatively characterized by the type of limitation (e.g. temperature or water) and quantitatively by the level of environmental stress. On the genetic side, the data showed qualitative differences induced by genotypes. Bustos-Korts et al. (2019a) and Bustos-Korts et al. (2019b) chose genotypic specifications of crop growth parameters that mimic realistic genetic variation for Australian environmental conditions.

### 2.4 Bayesian approach

Hierarchical models, of which our statistical model is a representative, lend themselves naturally to a Bayesian approach. Moreover, in our experience with hierarchical models, modern Bayesian software such as **Stan** is less prone to numerical failure than the off-the-shelf nonlinear least squares methods in **R** that were used by Van Voorn et al. (2023). See Comets et al. (2017), Driver and Voelkle (2018) and Plan et al. (2012) for a detailed discussion on algorithmic convergence issues in this context.

A data generating mechanism in a Bayesian model is described through the likelihood function. In line with typical assumptions of additive and normally distributed measurement errors, we assumed that the likelihood function was Gaussian and centered on values *M* (*t*_*k*_) at observation points *t*_*k*_, and had a constant standard deviation *σ*.

Another mandatory ingredient of Bayesian approach is a prior distribution on the model parameters. This is a probability distribution that reflects one’s prior knowledge on model parameters. It can also be viewed as a penalty or regularization term in the estimation procedure. Transtrum et al. (2011) explain the usefulness of priors as penalty terms in the context of non-linear least squares estimation. Our exact prior specifications will be given later on.

Except for simple statistical models, the posterior distribution is rarely available in analytic form. To address this issue, over the years powerful Markov Chain Monte Carlo (MCMC) methods have been developed to approximate posterior distribution through random sampling; see Brooks et al. (2011) for an overview. In general, MCMC methods are regarded as the gold standard for approximate posterior inference in that, in a long run, they provide asymptotically exact posterior samples. Their accuracy is offset by constraints of scalability and computational feasibility. Other options to approximate the posterior include variational inference (Blei et al., 2017), sequential Monte Carlo (Chopin and Papaspiliopoulos, 2020), Approximate Bayesian Computation (ABC) (Sisson et al., 2019), and the integrated nested Laplace approximation (INLA) (Gómez-Rubio, 2020).

### 2.5 Bayesian hierarchical model

The log-logistic model (4) was used to develop a hierarchical Bayesian model (Gelman and Hill, 2006; Gelman et al., 2014) for the APSIM wheat data from Section 2.3. The underlying idea of hierarchical modeling is that even though genotypes differ from each other, they have common features, and one can attempt to exploit this when fitting a model to the data. A similar idea applies to environmental variations over multiple years. As a result, data are used more efficiently, which, in principle, should lead to more accurate parameter estimates and a better quantification of the remaining uncertainty in inferential conclusions. This becomes particularly important in the real data setting where biomass measurements are obtained through relatively infrequent drone flights; see Chen et al. (2022), Smith et al. (2021), Panday et al. (2020a), Panday et al. (2020b) for an overview of longitudinal drone imagery.

In a nutshell, we assume that dynamics of crop growth in each trial and for each genotype can be described by a log-logistic function with an own set of parameters, whereas the remaining variation is attributed to statistical noise. The three parameters of the log-logistic function are themselves driven by genetic and environmental information. In the next step, we assume that parameters across trials and genotypes are linked through an overarching distribution. Inference is to be conducted on this latter distribution. The resulting statistical model is dynamic, but embedded within the mixed modeling framework at the parameter level (Pinheiro and Bates, 2000; Lee, 2022).

For the model description below, we provide a Kruschke-style diagram, cf. Kruschke (2014), in Figure 1. We use the following notation throughout: *y*_*ijk*_ is the biomass measurement at time *t*_*k*_ for genotype *i* and trial *j*. Next, *M*_*ijk*_ = *M* (*t*_*k*_; *θ*_*ij*_) is the value of the log-logistic function at time *t*_*k*_ for genotype *i* and trial *j*. The parameter *θ*_*ij*_ is a three-dimensional vector, its coordinates being logarithms of the maximum or right asymptotic biomass 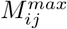, midpoint time 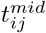 and growth rate *r*_*ij*_, respectively (for economy of notation, here we have redefined parameters *θ*_*ij*_ from Section 2.1 to their logarithms). Finally, *σ* is the measurement noise level.

**Figure 1:**
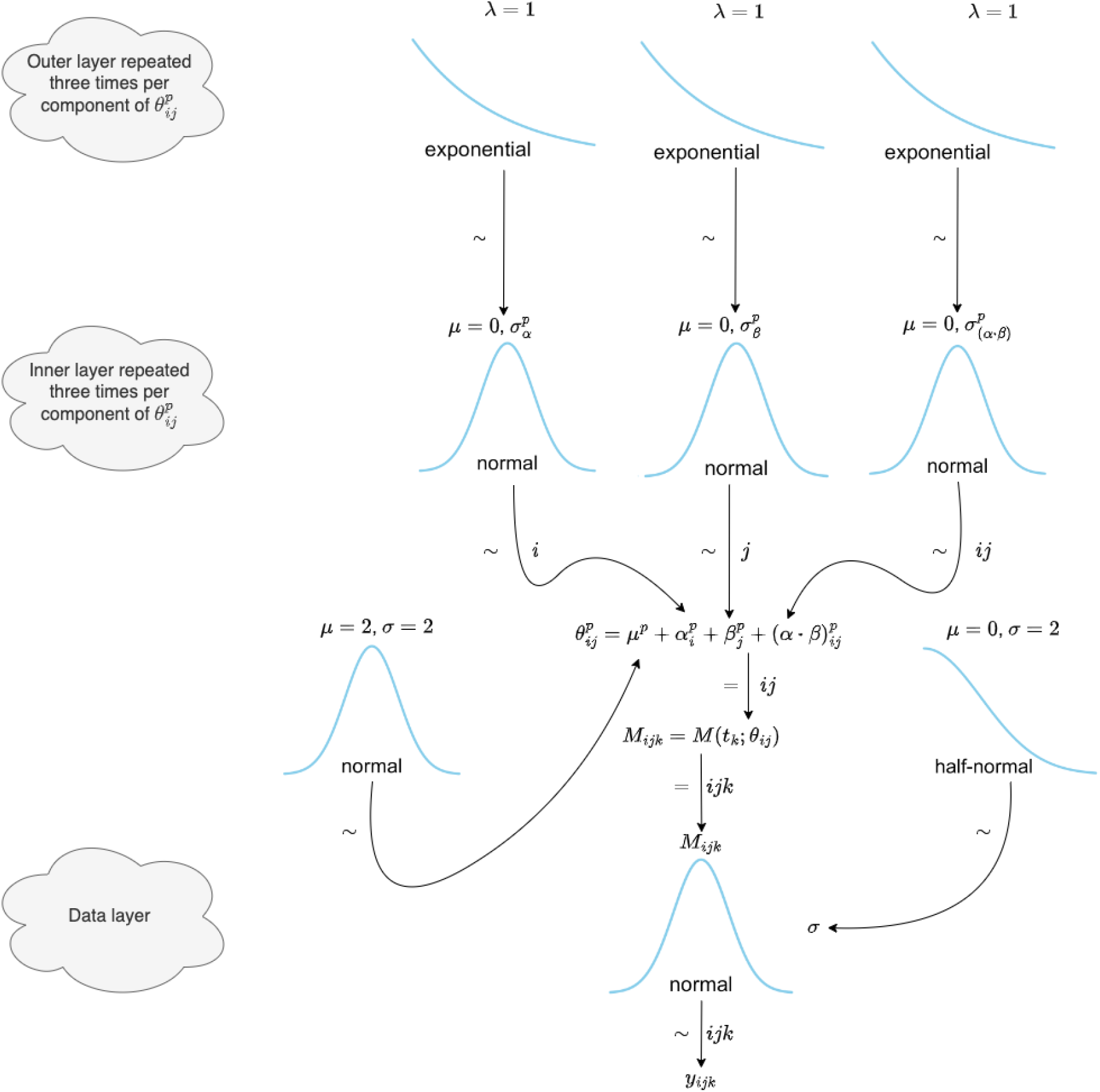
The Kruschke style diagram for the hierarchical Bayesian model from Section 2.5. See the main text for the notation. The inner and outer layers actually consist of three identical copies of distributions, although only one is shown. For each coordinate 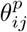 of the parameter vector *θ*_*ij*_, with the superscript varying over *p* = *M*, *mid* and *r*, a separate distribution is specified. The indices halfway the arrows indicate the way in which the variables or parameters are replicated over those indices.

As the construction is the same for each of the parameters of the log-logistic function, we focus on the right asymptote *M*^*max*^ in our presentation. Our key assumption is the decomposition

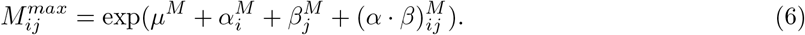

The exponentiation is to ensure positivity of the parameter, whereas working on the logarithmic scale enables one to build into it a two-way analysis of variance or ANOVA model structure

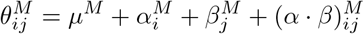

with, obviously, 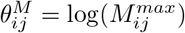. Next, *µ*^*M*^ can be thought of as the intercept common to all genotypes and trials. The term 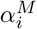 is contribution of the genotype *i* (on the logarithmic scale) to the DE parameter 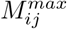. Similarly, 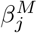 models the effect of specific environmental and management conditions in the trial *j* on the DE parameter 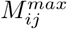. Finally, 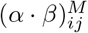 is the interaction term for genotype *i* and trial *j*. We treat 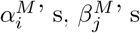 and 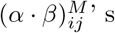 as random effects; see, for example, Pinheiro and Bates (2000). The linear model formulation for the logarithm of 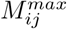 is an example of a GLMM (Searle et al., 2006; Faraway, 2016) for a DE parameter embedded in our hierarchical Bayesian framework for multi-genotype multi-environment dynamic data. This GLMM-like embedding offers a very powerful and versatile modeling framework with options for quantification of genotypic and environmental contributions to dynamic trait behavior.

The interpretability of the parameters of the log-logistic function can be directly leveraged to set priors in the Bayesian model. The intercept *µ*^*M*^ is assigned the prior

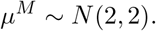

This prior is weakly informative: the distribution covers a likely range of the logarithm of the biomass measured at the end of the growing season for the APSIM data, while still allowing for a significant variation.

The genotype random effects 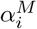 are assigned a prior through a hierarchical construction:

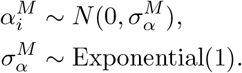

Similarly, for the trial random effects 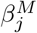 we assume

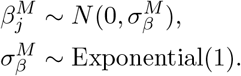

Also, these priors are weakly informative: the exponential distribution Exponential(1) has mean equal to 1. As a result, conditional on the values of 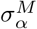 and 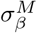, the parameters 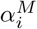 and 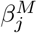 are assigned a broad prior, which nevertheless has a regularizing effect.

Finally, for the interaction random effect terms we assume

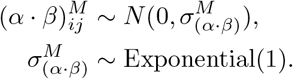

Priors used for parameters 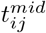 and *r*_*ij*_ can be found in the Supplementary Material accompanying the article. Like for 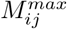, the key assumption is a two-way ANOVA structure

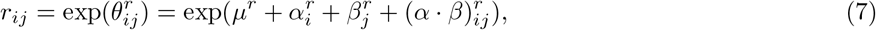

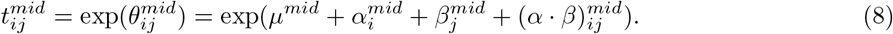

Interpretation of the terms in these formulae is analogous to the one in (6).

For the measurement noise level we use the Half-Normal prior

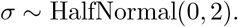

As the measurement errors *ε*_*ijk*_’s are assumed to be additive and independent Gaussian, this leads to the observation equation

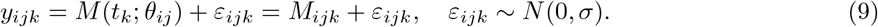

Thus for a genotype *i* and a trial *j*, the *y*_*ijk*_’s are the longitudinal growth data. The model in (6), (7), (8) and (9) is a GLMM with a non-standard link function for each of the DE parameters. The non-standard link follows from a composition of the exponential function and the ODE solution (log-logistic function). Normality of measurement errors is a reasonable assumption as a first approximation in the present setting. In fact, daily APSIM wheat data were generated without noise, and therefore the unexplained residual variation can be expected to be relatively small. The error standard deviation *σ* quantifies the lack of fit of the present model. The wide spacing of the data in time is expected to mitigate correlations between residuals. See also Section 2.2 in Seber and Wild (1989).

Priors in Bayesian analyses can be set based on the experience of the user, knowledge of the problem at hand, convenience, and regularization or numerical stability considerations. For data from plant breeding experiments, the variation between trials is typically much larger than that between genotypes, while *G* × *E* is again smaller than variation between genotypes. We did not implement that knowledge in the current analyses and used only weakly informative priors for our DE parameters, because we knew that the data would be highly informative, while the prior effects would be limited. Nowadays, many Bayesians are comfortable with using data-dependent priors; see, e.g., Section 9.5 in Gelman et al. (2021) for the rationale behind default priors in the popular **R** package **rstanarm**, which is similar to the **brms** procedure we used for our analyses in this paper, and cf. Gelman et al. (2017). The abundance of data (specifically, the data for genotypes outside our subset of 48 genotypes) in our case offered some possibilities for fine-tuning and experimentation with priors: for instance, we inspected where the observed growth curves level off and centered the priors so that this range receives substantial prior mass. As a qualification, we remark that we aimed at investigating dynamic phenotypes at medium-time resolution, i.e., biweekly, as can occur in experiments with regular drone flights. The number of observations per growing season for such experiments is maximally around 10 to 15. This number restricts the complexity of the DE system that can be successfully fitted to such data.

### 2.6 Parameter identifiability

Although in Section 2.5 the logarithms of the parameters of the differential equation, i.e. 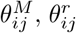 and 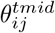, are identifiable, their additive decompositions into the sums of the main genotypic and environmental effects and their interactions are not unique. This is well-known for ANOVA-type models. Additional restrictions must be imposed to talk about genotypic, environmental, and *G* × *E* effects. Thus, once the MCMC samples of the parameters have been collected, in the post-processing step we applied the sum-to-zero restrictions on 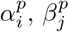 and 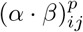, with *p* being one of *M*, *t*^*mid*^ and *r*. See, e.g., Kruschke (2014), Section 20.1.1, Gelman and Hill (2006), Section 19.4, and Ogle and Barber (2020) for details. Main effects and interactions refer to parameters in this new parametrization, and are unambiguously defined. In principle, the new parameters need to receive other names, such as 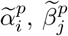 and 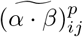. However, for notational simplicity, in the rest of the paper we will keep the old notation 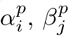 and 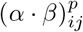, but we will mean the parameters in the new parameterization. We also remark that priors in Bayesian mixed models help with weak identifiability of parameters. In fact, the conclusions in this paper would have stayed largely unchanged, had we used in our analysis the raw random effects instead of the adjusted ones.

### 2.7 Interpretation of genotypic, environmental and *G* × *E* parameters

Interpretations of dynamic behavior shown by individual genotypes and induced by individual environments can be based on estimates for individual effects and their dispersion parameters building on the interpretation of similar parameters in traditional quantitative genetic models for continuous traits. To be more specific, we will build such interpretations on the posterior distributions for genotypic and environmental main effects and *G* × *E* effects, i.e., 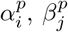 and 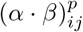, respectively, with *p* being one of *M*, *t*^*mid*^ and *r*, together with the corresponding dispersion parameters 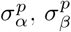, and 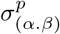. Inspection of the posterior distributions of the dispersion parameters will inform us about the relative importance of genotypic and environmental main effects as well as that of *G* × *E*. To appreciate the relative magnitudes of *G* × *E* effects versus genotypic main effects for the dynamic parameters, we will look at the posteriors for the ratios of the *G* × *E* standard deviations to the standard deviations for genotypic main effects.

For dynamic parameters with sizeable dispersion for *G* × *E*, individual interaction effects, 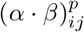, can be studied via their credible intervals to verify whether they contain zero. Questions concerning genotypic and environmental characterizations can be addressed by counts on non-null *G* × *E* effects. Genotypes can be characterized on the percentage of times they show non-null *G* × *E* effects across all trials or subsets of trials. This percentage can be interpreted as a measure of stability. The four locations that were included in the simulation data represent different growing regions within Australia. Therefore, it may be of interest to see to which extent the incidence of non-null *G* × *E* effects differs between those regions. A low incidence may point to a non-discriminative environment, whereas a high incidence would point to a discriminative environment.

Posterior distributions of environmental and/or *G* × *E* effects can be used for environmental characterization to create so-called environmental scenarios or envirotypes (Xu, 2016; Resende et al., 2021; Chen et al., 2026; Fernandez et al., 2026). Typically in environmental characterization, trials are clustered into groups, with the groups representing a distinguishable type of growing environment. Envirotypes can serve to develop more efficient breeding strategies. We illustrate how to use posterior medians for the environmental main effect parameters 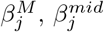, and 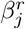 as inputs for an environmental characterization.

These posterior medians were standardized and collected in a vector of length three. These vectors were clustered into three groups using a *K*-means algorithm (MacQueen, 1967). Choosing three or four clusters is supported by relevant literature on envirotypes (Fernandez et al., 2026). The results of the *K*-means clustering depend on the random initialization of the algorithm. With three groups, the inferred clusters were stable over multiple runs of the algorithm.

For plotting purposes, we used for some figures medoid representations (Van der Laan et al., 2003). For environments, this resulted in real environments that were at the center of the space spanned by estimates for the environmental main effects in the GLMMs for the three parameters. The medoid environment can then be thought of as a typical environment within a class of environments. The same technique was also used for plotting genotypic information.

### 2.8 Software implementation

We implemented our Bayesian approach in **Stan**, specifically using the **R** package **brms. Stan** is a state-of-the-art platform for statistical modeling and high-performance statistical computation. The principal engine to perform Bayesian inference in **Stan** is the No-U-Turn-Sampler (NUTS), a cutting-edge variant of the Hamiltonian Monte Carlo (Hoffman and Gelman, 2014; Betancourt, 2017; Betancourt et al., 2017). The **R** package **brms** uses **Stan** as its backend, but employs the formula syntax similar to the popular **lme4** package. This results in a familiar and simple interface for performing regression type analyses. See Bürkner (2017, 2018, 2021) for additional information. For a general overview and comparison of the modern software for Bayesian inference, see Štrumbelj et al. (2024).

**brms** automatically evaluates the popular 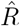 convergence diagnostic for Markov chains it generates (Gelman and Rubin, 1992; Vehtari et al., 2021), and issues a warning when its values are above the commonly used threshold 1.05. See also Table 5 in the Supplementary Material for values of 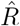, which are all below the threshold 1.05. In order to verify convergence, we also visually examined the trace plots and autocorrelation functions for our Markov chains to the extent that this is possible given the large number of parameters in the model (Gabry et al., 2025). The plots for an arbitrary selection of the model parameters are presented in the Supplementary Material in Figures 11 and 12. From Table 5 we also see that the lowest effective sample sizes occur for the global intercept parameters, which are weakly identified from the data and are not of inferential interest. On the other hand, the standard deviation parameters have bulk and tail effective sample sizes above 400, which allows for accurate simulation-based inference in the present context.

To handle the **brms** output, we mainly employed the **R** package **tidybayes** (Kay, 2022), and other capabilities offered by **R** (R Core Team, 2022). For plotting in **R** we relied on **ggplot2** (Wickham, 2016), as well as several other packages building upon and extending **ggplot2**.

In our implementation in **brms**, we used an explicit form of the log-logistic function given in (4). For future extensions, we note that availability of an analytic solution to a differential equation is not a requirement for our Bayesian approach: it also works when the analytic solution is replaced with a numerical solution. In that case, instead of **brms**, the model may be implemented directly in **Stan**; cf. Chatzilena et al. (2019) and Grinsztajn et al. (2021). Alternatively, additional **Stan** code can be injected into the **brms** model specification. Fitting, however, is faster with an analytic solution. As for the effect of the approximation error on posterior inference resulting from numerical integration, the impact stays bounded, and is negligible when the step size of the ODE solver is chosen to be small enough; see Donnet and Samson (2007) for a mathematically precise statement. **Stan** implements a number of reliable adaptive ODE solvers (Stan Development Team, 2023, Chapter 13). For early contributions to Bayesian fitting of hierarchical differential equation models, see Bois et al. (1996); Gelman et al. (1996); Wakefield (1996). These early contributions rely on Gibbs sampling (Geman and Geman, 1984; Gelfand and Smith, 1990). However, the Gibbs sampler is known to suffer from poor mixing in statistical problems with highly correlated parameters in the posterior (De Jong and Shephard, 1995; Lindsten and Schön, 2013). This is inherent in the sampler because of its sequential updating of single individual parameters in any given iteration. This slow convergence hampers the scalability of the Gibbs sampler to high-dimensional models with complex posterior geometry, such as those defined by differential equations (Girolami, 2008; Calderhead and Girolami, 2009), and especially their hierarchical extensions.

## 3 Results

### 3.1 Goodness of fit and characterization of sources of variation

To assess how well our model fitted the ‘observed’ longitudinal data for individual genotypes in individual environments, we looked at plots of observed (Figure 2) and fitted (Figure 3) data. The observations were created by sampling at medium-time resolution biomass data from APSIM wheat simulations. There were four locations with 31 years of trials, in total 124 environments, for a set of 48 genotypes. In Figure 2, for each location we present the so-called medoid trial, a trial that can be thought of as best representing the group (location) to which it belongs. For such a medoid trial, we show the daily data for all genotypes. The colors of the genotypes are consistent in Figures 2 and 3. Moreover, to give an idea of the amount of uncertainty in the obtained fits, in Figure 4 we show 95% Bayesian pointwise prediction bands for the medoid genotype in the medoid year (trial) at each of the 4 locations. As can be readily seen, most observations fall within the prediction bands.

**Figure 2:**
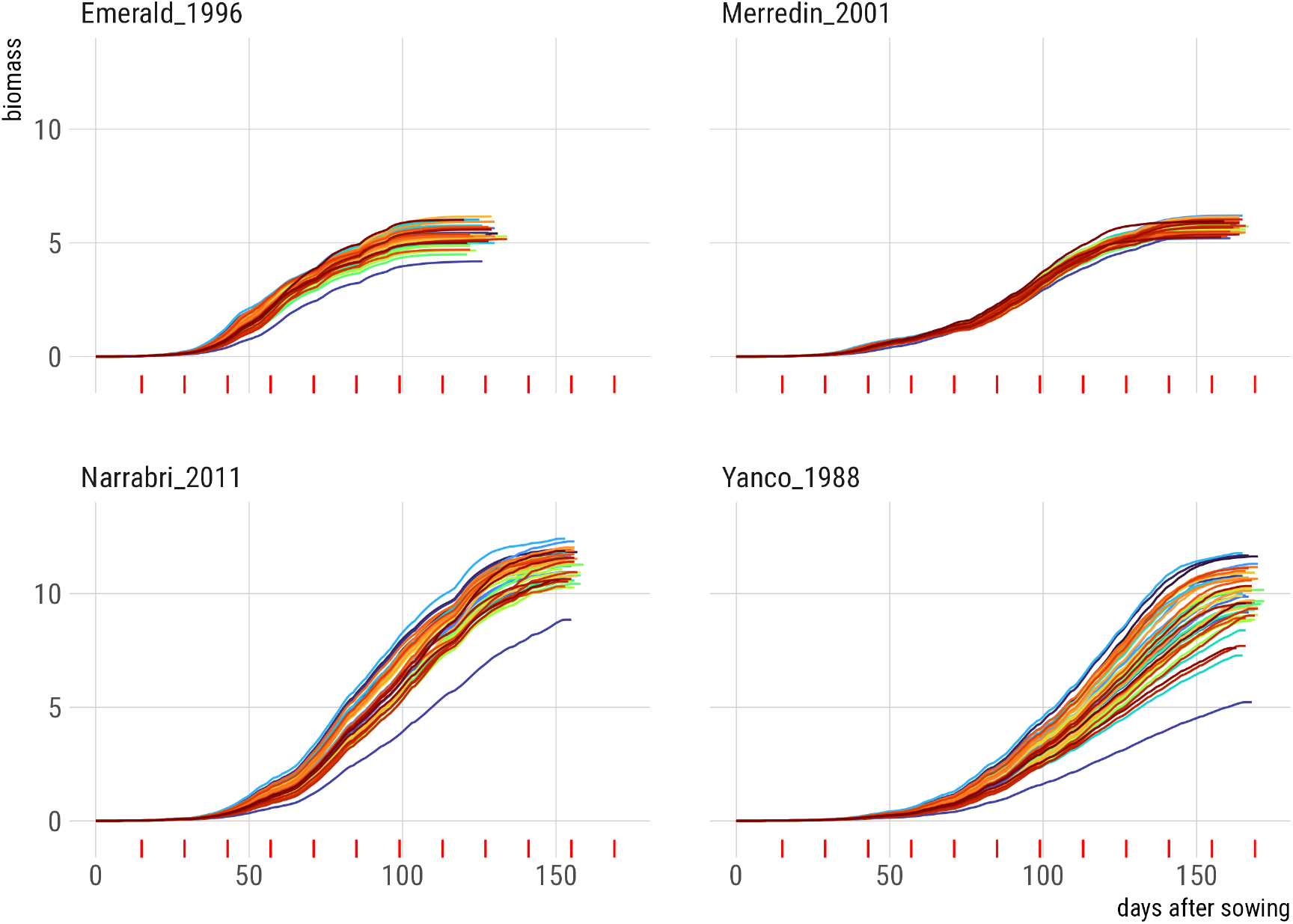
Observed daily APSIM wheat biomass data for medoid year at each location showing all 48 genotypes. Lines are colored according to genotypes. Colors are consistent across different panels. The red tick marks along the horizontal axis indicate time points at which biomass observations were used for Bayesian hierarchical modeling.

**Figure 3:**
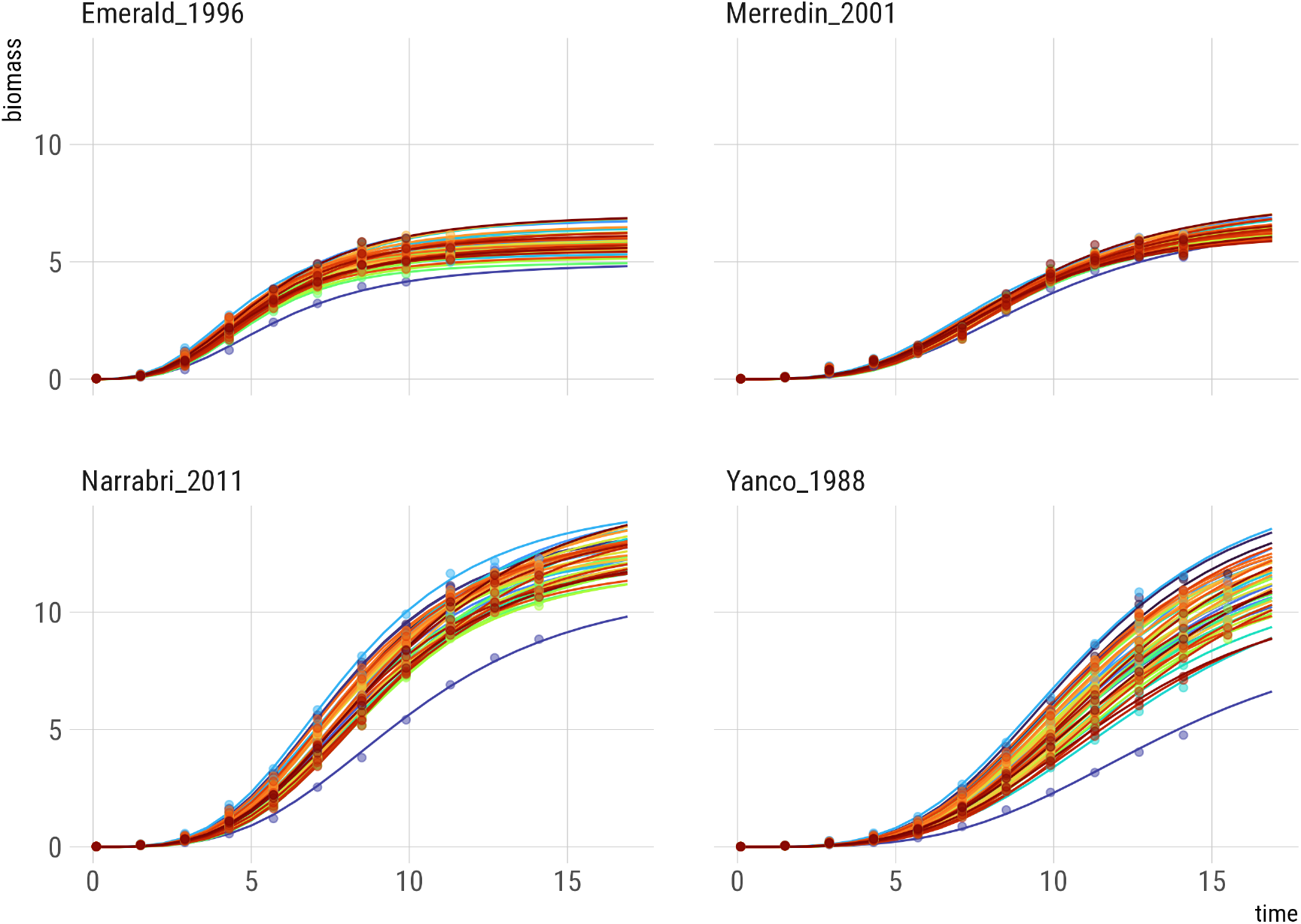
Fitted curves (posterior medians) for medoid year at each location showing all genotypes. Points represent observations. Lines and points are colored according to genotypes. Colors are consistent across panels. The time axis has been obtained by dropping the first 14 days after sowing and rescaling by division by 10.

**Figure 4:**
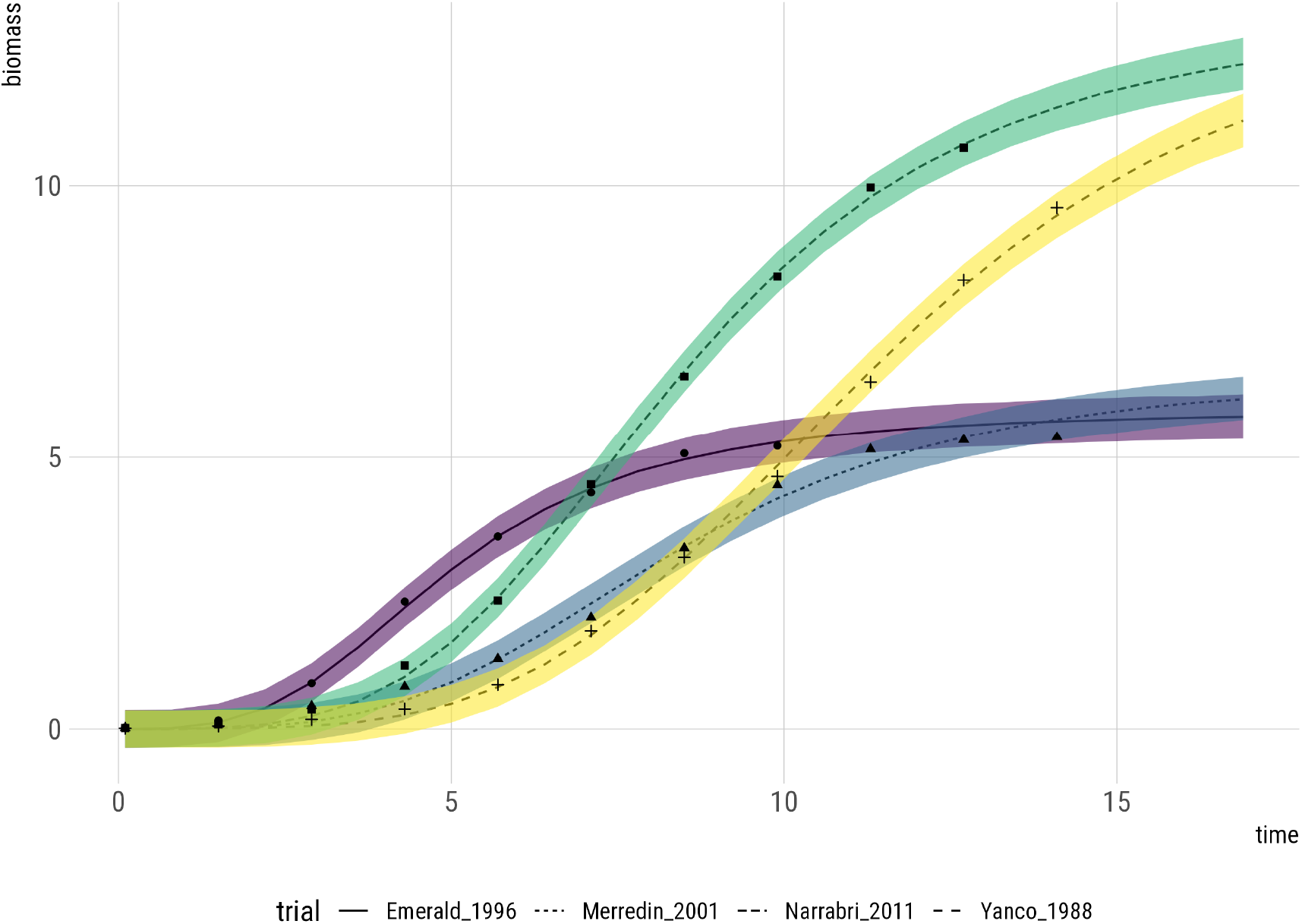
Fitted curves (posterior medians) for medoid genotype in medoid year for each of 4 locations. Observations at locations are distinguished by symbol: points, squares, triangles and crosses. Colored bands represent 95% Bayesian (pointwise) prediction bands. Bands are colored according to locations. The time axis has been obtained by dropping the first 14 days after sowing and rescaling by division by 10.

The log-logistic fits were good in that they follow the observed data closely for all genotypes and trials, and for most predictive purposes in this setup the log-logistic fits are probably good enough. Still, the observed data contain small bumps, deviations from the smooth log-logistic fits, which are very likely caused by short-term meteorological events, such as elevated temperatures and drought. Such bumps cannot be incorporated into the log-logistic model, but can show up in residual plots, see Figure 13.

Figure 2 contains genotypic curves that hardly intersect, but there are clear differences between locations, between years within locations (trials) and between genotypes. Narrabri and Yanco attained higher biomasses than Merredin and Emerald, with Emerald having the lowest biomasses. The amount of variation between years within locations seems comparable across the four locations, see Figure 5.

**Figure 5:**
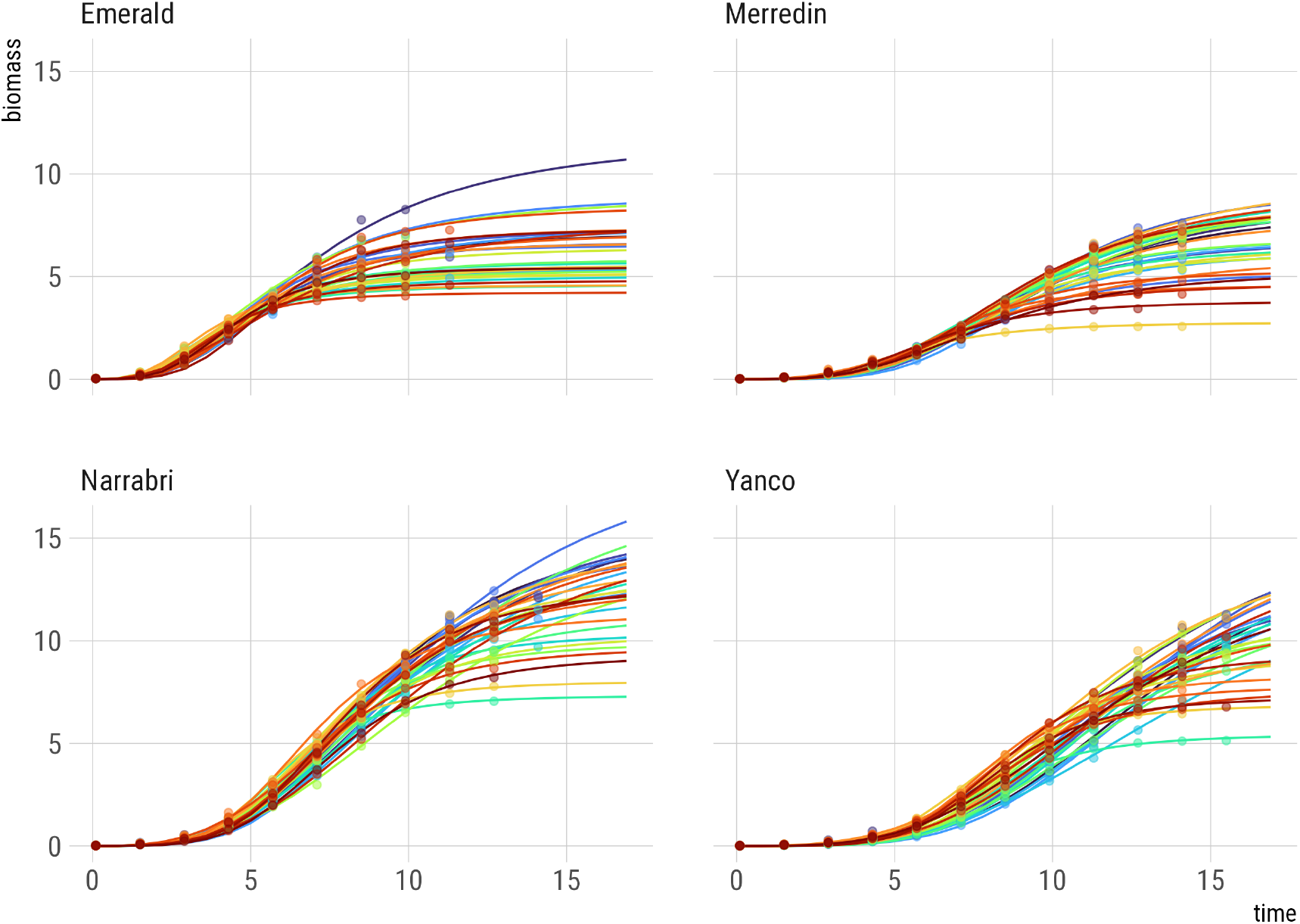
Fitted curves (posterior medians) for the medoid genotype across all years at each of 4 locations. Points represent observations. Lines and points are colored according to years. Colors are consistent across panels. The time axis has been obtained by dropping the first 14 days after sowing and rescaling by division by 10.

Inspection of the posterior standard deviations for environmental and genotypic main effects and *G* × *E* effects in Figure 6 shows that the dynamic parameters for asymptote, midpoint and slope follow the standard partitioning of phenotypic variation with environmental main effects being responsible for the majority of the phenotypic variation, followed by genotypic main effects and finally *G* × *E* effects. The greatest variation was found in the asymptote, followed by the midpoint, while the slope was clearly less variable in this wheat panel. *G* ×*E* variation is best appreciated in relation to genotypic main effect variation. Figure 7 presents ratios of *G* × *E* standard deviation to genotype standard deviation. For the asymptote, this ratio was between 0.6 and 0.9, while for the midpoint it was between 0.25 and 0.35. For the slope, the same ratio had a value between 0.0 and 0.1. The *G* × *E* appeared clearly for the asymptote, but for the midpoint and slope parameter it had little to no influence. Therefore, the rough biomass growing pattern for wheat seems predictable from additive GLMMs for at least midpoint and slope. That is, no or hardly any *G* × *E* plays a role for midpoint and slope parameters in the biomass growth functions. For the asymptote, the maximum biomass, *G* × *E* does play a role, and further study is required to find out how to arrive at a predictive model for biomass taking into account the *G* × *E* for maximum biomass.

**Figure 6:**
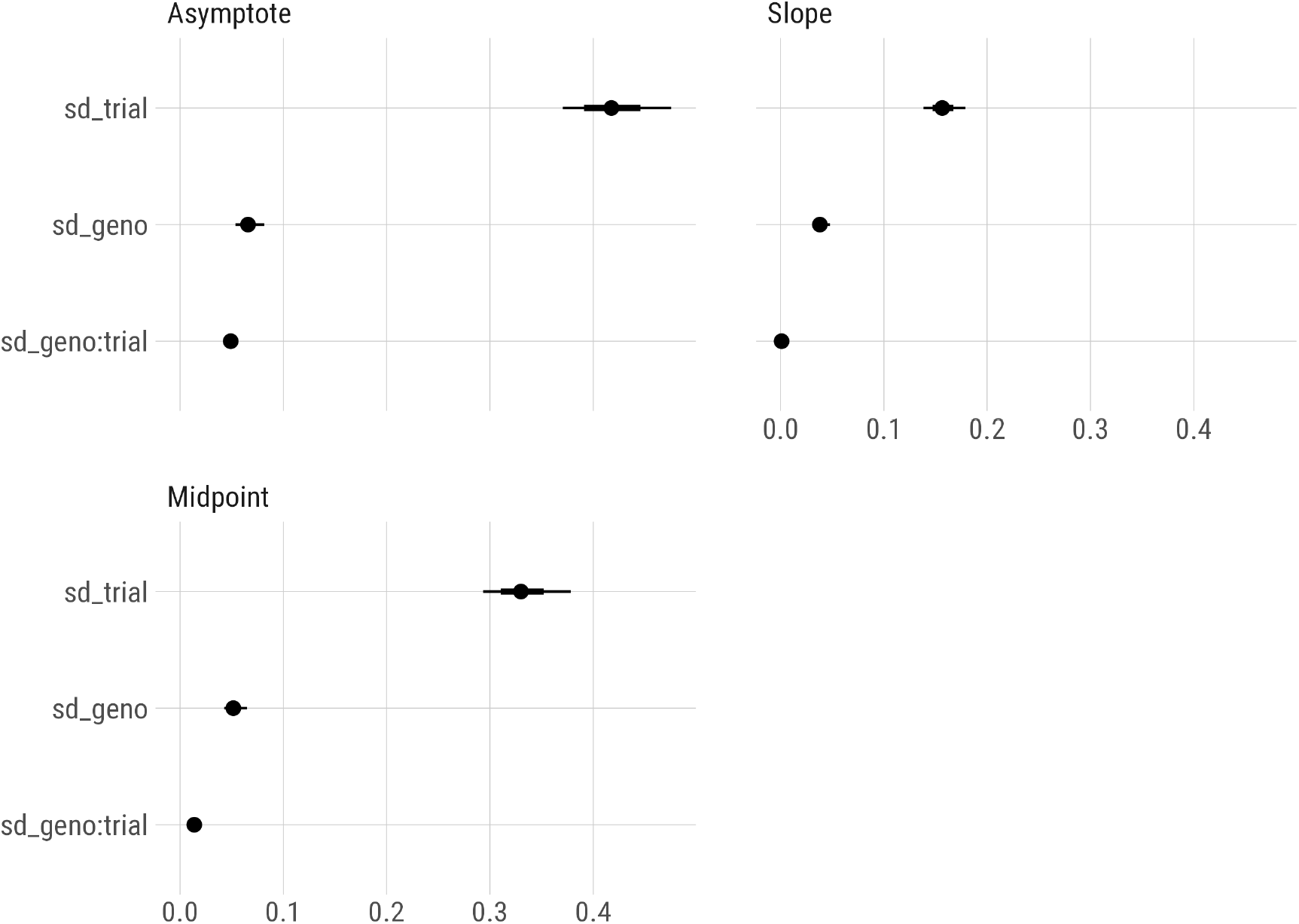
Posterior summaries for parameters 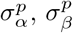, and 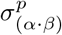 for each of the superscripts *p* = *M*, *mid* and *r*. Each panel gathers results for a different parameter of the log-logistic function, identified by labels `Asymptote`, `Midpoint` and `Slope`. Points in the middle of the segments are posterior medians. The thick and thin segments represent 67% and 95% credible intervals, respectively. No posterior densities are plotted, as they are highly peaked for 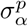 and 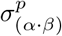 parameters, and as such visually uninformative.

**Figure 7:**
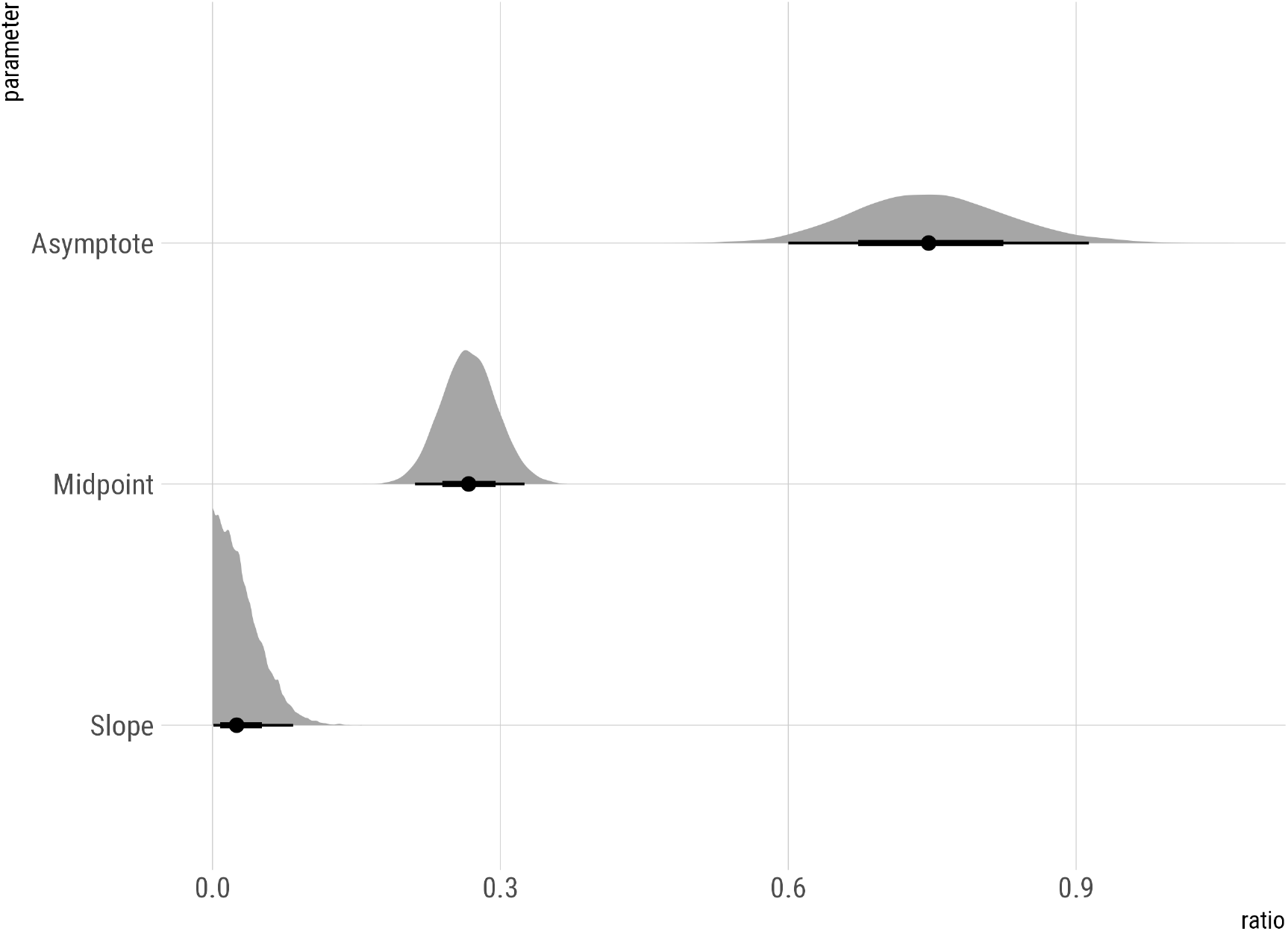
Posterior summaries for ratios 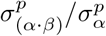 for each of the superscripts *p* = *M*, *mid* and *r*. Labels `Asymptote`, `Midpoint` and `Slope` refer to the parameters of the log-logistic function. Points in the middle of the segments are posterior medians. The thick and thin segments represent 67% and 95% credible intervals, respectively. The densities are the posterior densities.

### 3.2 G × *E* interactions

Interactions can be further studied via credible intervals for interaction terms 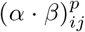, and verifying whether zero is contained within these intervals or not. Table 2 summarizes the results and confirms the impression that with the log-logistic model, most of the interaction is to be discovered for the asymptote, 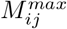. For this DE parameter, *G* × *E* did occur in about a third of all *G* × *E* combinations across the full set of trials. More detail is provided by the number or percentage of *G* × *E* interactions per location in Table 3. For parameters 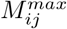, there is some variation between locations in the frequencies of interactions, with *G × E* being more frequent in Yanco and less frequent in Merredin. Further zooming on *G* × *E* interactions, we inspect the non-zero interactions in the trials over time (years) for each of the locations. In Figure 8, we again see that Yanco in general induces more *G* × *E* than the other three locations, with Merredin inducing the lowest frequency of *G* × *E*. We also see that there is a lot of variation in the frequency of non-null *G* × *E* between years within locations.

**Table 1:**
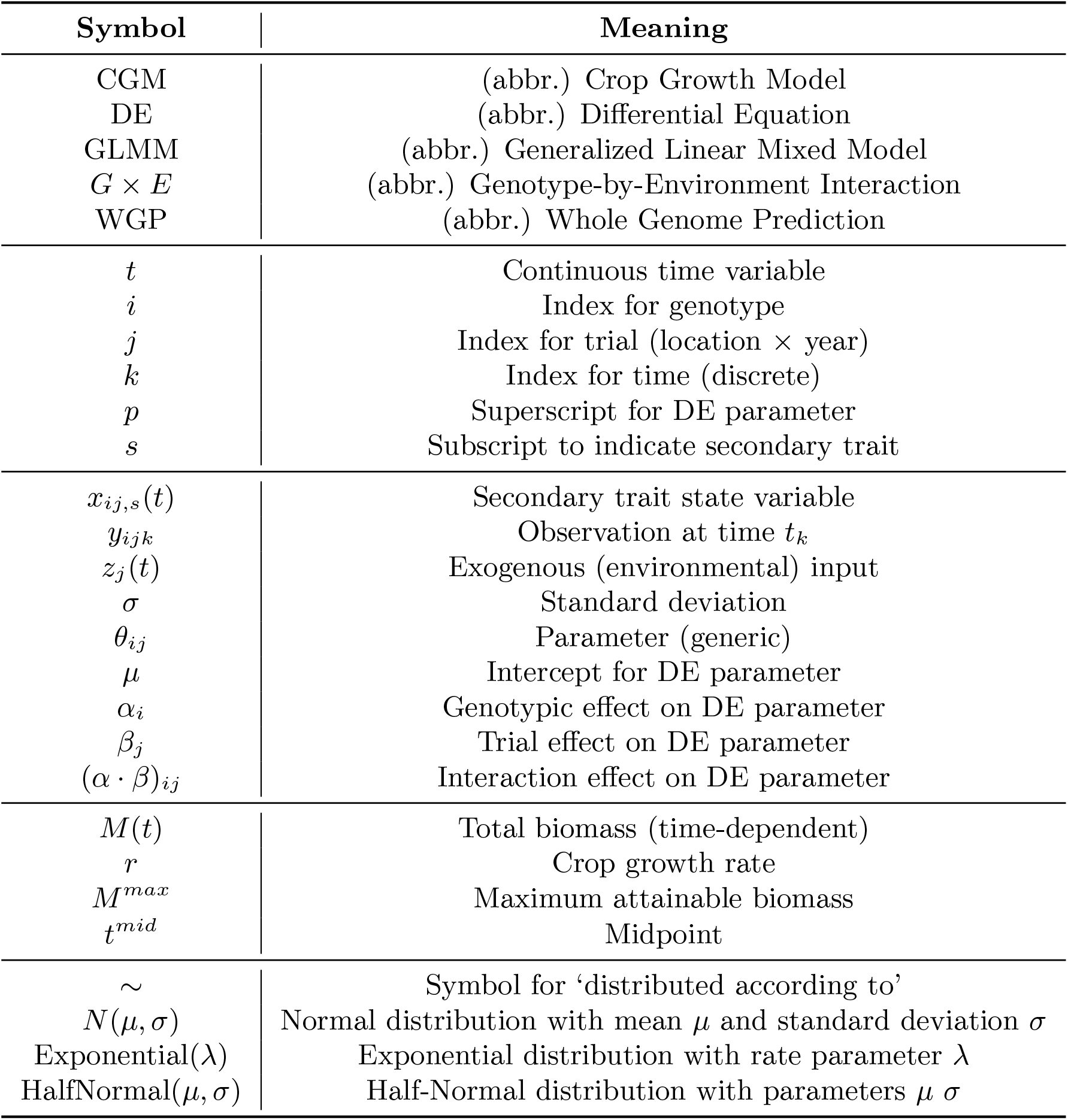
Explanations of the symbols and abbreviations used throughout this paper.

| **Symbol** | **Meaning** |
|---|---|
| CGM | (abbr.) Crop Growth Model |
| DE | (abbr.) Differential Equation |
| GLMM | (abbr.) Generalized Linear Mixed Model |
| *G* × *E* | (abbr.) Genotype-by-Environment Interaction |
| WGP | (abbr.) Whole Genome Prediction |
| *t* | Continuous time variable |
| *i* | Index for genotype |
| *j* | Index for trial (location × year) |
| *k* | Index for time (discrete) |
| *p* | Superscript for DE parameter |
| *s* | Subscript to indicate secondary trait |
| *x*<sub>ij,s</sub>(*t*) | Secondary trait state variable |
| *y*<sub>ijk</sub> | Observation at time *t*<sub>k</sub> |
| *z*<sub>j</sub>(*t*) | Exogenous (environmental) input |
| σ | Standard deviation |
| θ<sub>ij</sub> | Parameter (generic) |
| μ | Intercept for DE parameter |
| α<sub>i</sub> | Genotypic effect on DE parameter |
| β<sub>j</sub> | Trial effect on DE parameter |
| (α · β)<sub>ij</sub> | Interaction effect on DE parameter |
| *M*(*t*) | Total biomass (time-dependent) |
| *r* | Crop growth rate |
| *M*<sup>max</sup> | Maximum attainable biomass |
| *t*<sup>mid</sup> | Midpoint |
| ∼ | Symbol for 'distributed according to' |
| *N*(μ, σ) | Normal distribution with mean μ and standard deviation σ |
| Exponential(λ) | Exponential distribution with rate parameter λ |
| HalfNormal(μ, σ) | Half-Normal distribution with parameters μ σ |

**Table 2:** Frequencies of credible intervals for interaction terms 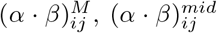 and 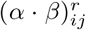 that exclude zero, i.e., that do indicate *G* × *E*.

|  | (α · β)<sub>ij</sub><sup>M</sup> | (α · β)<sub>ij</sub><sup>mid</sup> | (α · β)<sub>ij</sub><sup>r</sup> |
|---|---|---|---|
| Credible interval excludes zero | 2173 | 91 | 0 |
| Percentage of intervals without zero | 36.5% | 1.5% | 0% |

**Table 3:** Frequencies of credible intervals for interaction terms 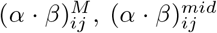 and 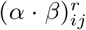 that exclude zero, i.e., that do indicate *G* × *E*. Frequencies are grouped by location.

|  | (α · β)<sub>ij</sub><sup>M</sup> | (α · β)<sub>ij</sub><sup>mid</sup> | (α · β)<sub>ij</sub><sup>r</sup> |
|---|---|---|---|
| Emerald | 534 | 0 | 0 |
| Merredin | 422 | 15 | 0 |
| Narrabri | 586 | 41 | 0 |
| Yanco | 631 | 35 | 0 |
| Total | 2173 | 91 | 0 |

**Figure 8:**
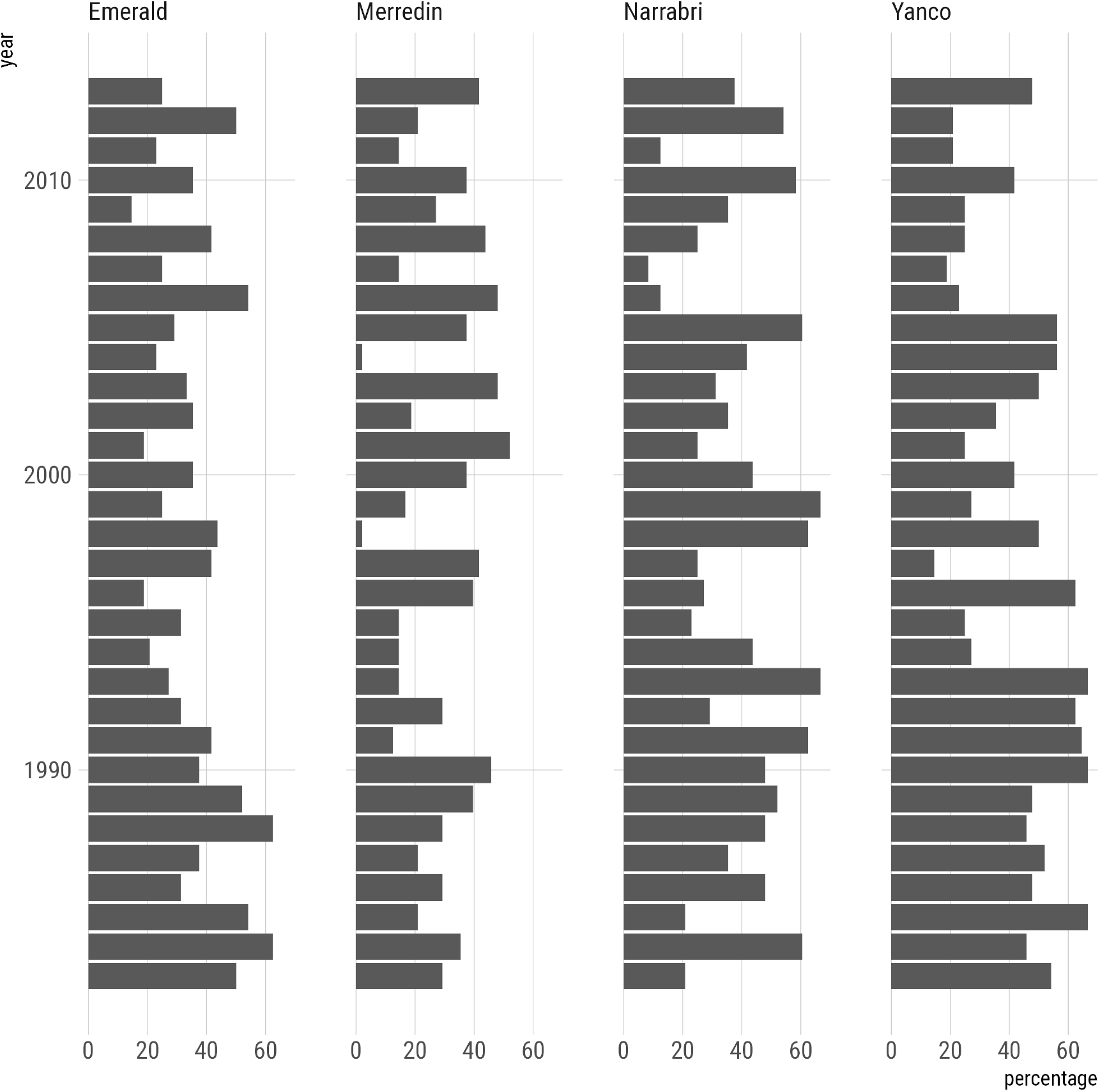
Percentages of genotypes in individual trials with interactions 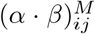 (for the asymptote parameter *M*^*max*^) whose 95% credible interval excludes zero. The higher this percentage, the more specific are the environmental challenges imposed by the conditions.

When we apply the same logic to the genotypes, we can compare genotypes for the number of non-null *G*× *E* effects across trials. We can interpret this number as an indicator for the genotypic stability of DE parameters. The lower the frequency or number of non-null interactions, the higher the stability of the genotype. Figure 9 offers a view on substantial genotypic variation for non-zero interactions for *G* × *E* in the asymptote, 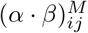.

**Figure 9:**
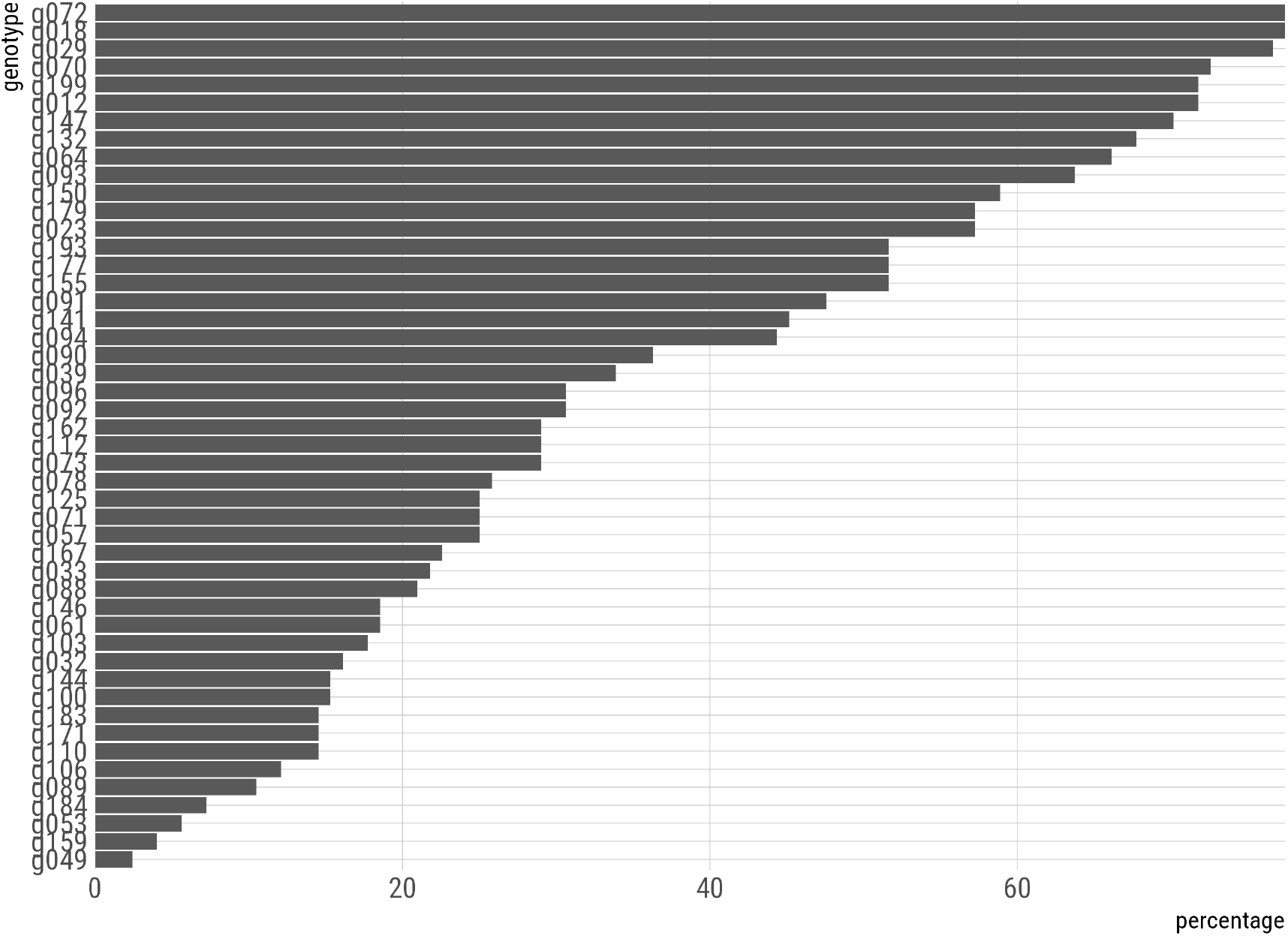
Percentage of interactions for individual genotypes 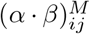 across all trials, whose 95% credible intervals exclude zero. This percentage can be read as a measure of stability. The lower the percentage, the more stable is a genotype.

### 3.3 Environmental characterization

The parameters in the GLMMs for the DE parameters can be used to characterize genotypes and environments. For the environments, the environmental main effect parameters 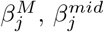, and 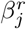 should reflect the local growing conditions in ways comparable to physical and physiological measurements. Upon collecting these parameters in three-dimensional vectors, we used the *K*-means clustering algorithm on these environmental descriptors. The *K*-means solution was stable for solutions with three clusters over multiple runs with different initializations. On inspection of these clusters, it turned out that Emerald and Merredin trials each dominated one cluster, while a third cluster was a combination of especially Narrabri and to a lesser extent Yanco trials. Envirotypes facilitate the efficient modeling of *G* × *E* by partitioning the initial *G* ×*E* in a part explained by *G × Envirotype* interaction and a residual part. For the APSIM wheat data here, because the envirotypes were dominated by differences between locations, a description of *G* × *E* for the asymptote by introducing a *G* × *Envirotype* term was about as effective as introducing a *G* × *Location* term, and a comparable residual *G* × *E* remained.

### 3.4 Genotypic characterization

In Bustos-Korts et al. (2019b), genotype-specific values for a panel of 199 genotypes were generated for 12 APSIM parameters that determine phenology, capture of environmental resources, resource use efficiency, and biomass partitioning. APSIM parameters were created from a genetic basis consisting of 300 QTLs whose effects were sampled from gamma distributions with parameters following estimates obtained from analysing real wheat data.

We wanted to characterize the genotypes in our hierarchical analysis by the estimated genotypic main effects in the GLMMs for the DE parameters. To this end, we compared the estimates for these genotypic effects with APSIM parameters used to generate the data by Bustos-Korts et al. (2019b). APSIM parameters that changed over time were not taken into account. The selected APSIM parameters are summarized in Table 4, which is constructed as a subset of Table 2 in Bustos-Korts et al. (2019b).

**Table 4:**
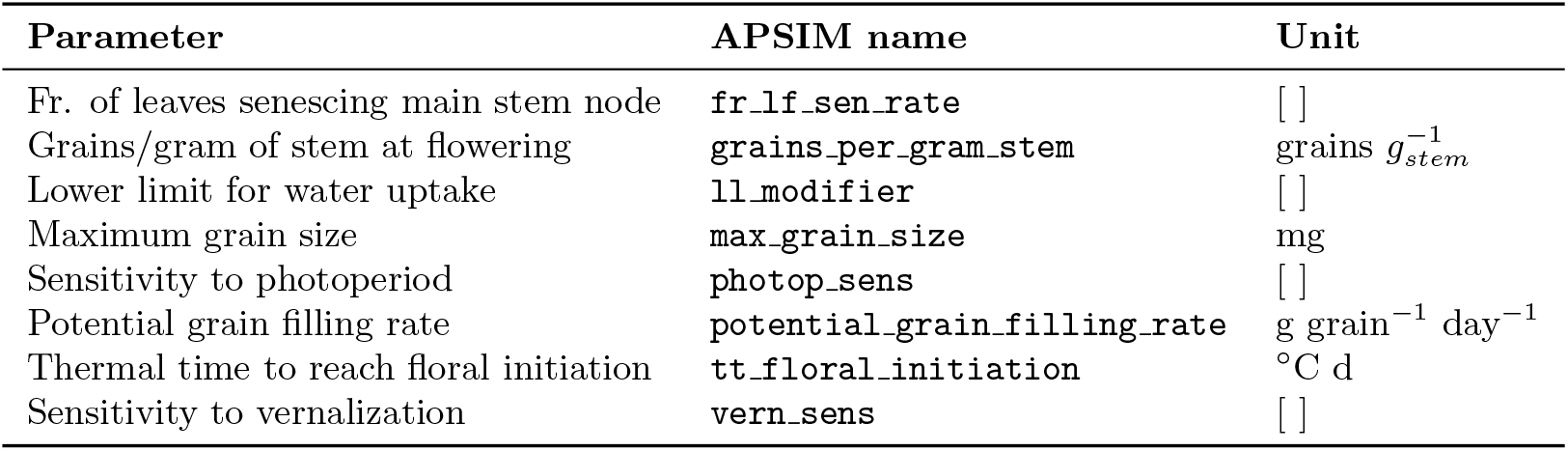
Genotype-specific parameters from APSIM that were used in this study for comparison purposes.

| **Parameter** | **APSIM name** | **Unit** |
|---|---|---|
| Fr. of leaves senescing main stem node | fr_lf_sen_rate | [ ] |
| Grains/gram of stem at flowering | grains_per_gram_stem | grains *g*<sub>*stem*</sub><sup>−1</sup> |
| Lower limit for water uptake | ll_modifier | [ ] |
| Maximum grain size | max_grain_size | mg |
| Sensitivity to photoperiod | photop_sens | [ ] |
| Potential grain filling rate | potential_grain_filling_rate | g grain<sup>−1</sup> day<sup>−1</sup> |
| Thermal time to reach floral initiation | tt_floral_initiation | °C d |
| Sensitivity to vernalization | vern_sens | [ ] |

**Table 5:**
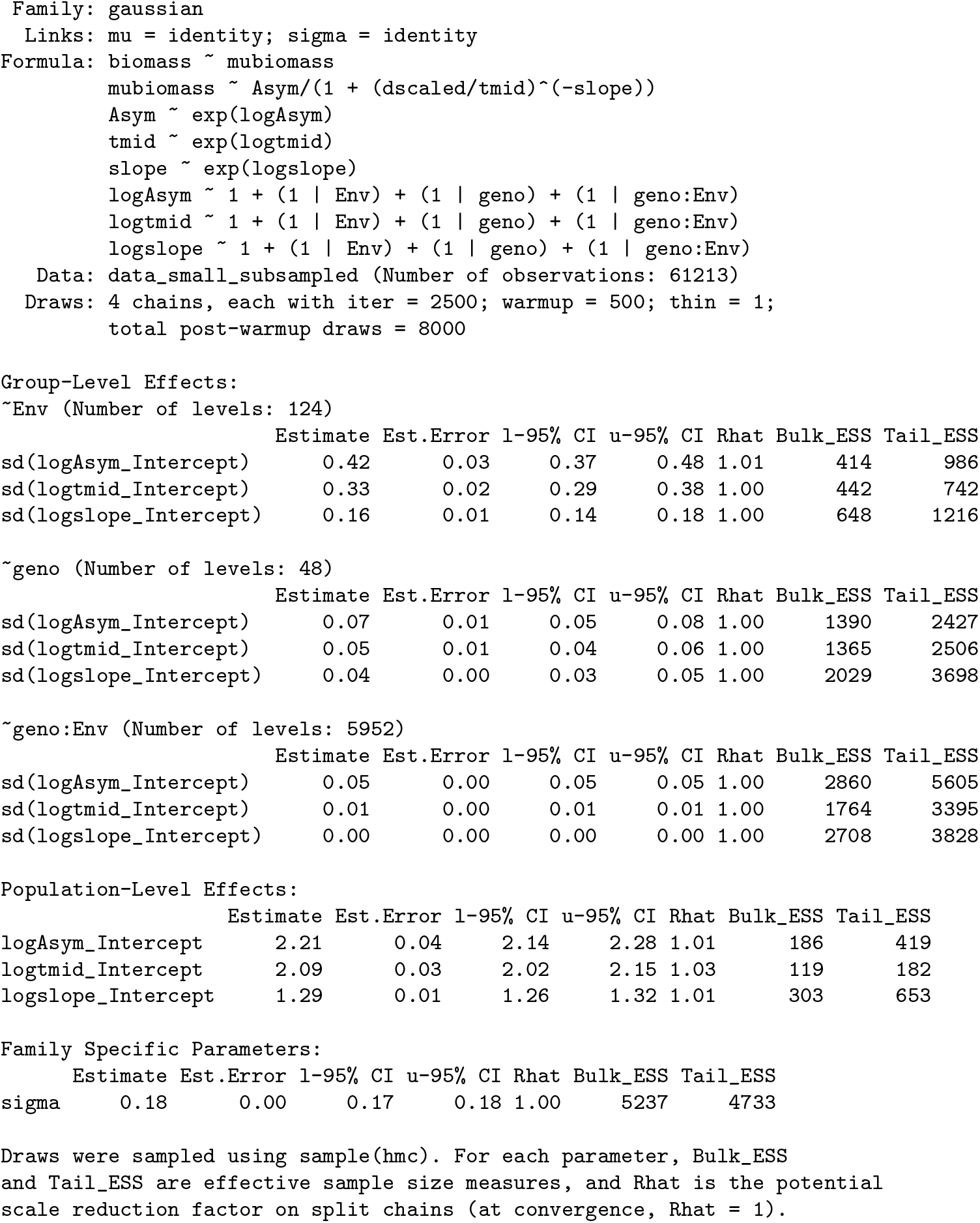
Summary of the brms fit.

|  | Estimate | Est.Error | l-95% CI | u-95% CI | Rhat | Bulk_ESS | Tail_ESS |
|---|---|---|---|---|---|---|---|
| Family: gaussian |  |  |  |  |  |  |  |
| Links: mu = identity; sigma = identity |  |  |  |  |  |  |  |
| Formula: biomass ~ mubiomass<br>mubiomass ~ Asym/(1 + (dscaled/tmid)^(-slope))<br>Asym ~ exp(logAsym)<br>tmid ~ exp(logtmid)<br>slope ~ exp(logslope)<br>logAsym ~ 1 + (1 \| Env) + (1 \| geno) + (1 \| geno:Env)<br>logtmid ~ 1 + (1 \| Env) + (1 \| geno) + (1 \| geno:Env)<br>logslope ~ 1 + (1 \| Env) + (1 \| geno) + (1 \| geno:Env) |  |  |  |  |  |  |  |
| Data: data_small_subsampled (Number of observations: 61213) |  |  |  |  |  |  |  |
| Draws: 4 chains, each with iter = 2500; warmup = 500; thin = 1;<br>total post-warmup draws = 8000 |  |  |  |  |  |  |  |
| Group-Level Effects: |  |  |  |  |  |  |  |
| ~Env (Number of levels: 124) |  |  |  |  |  |  |  |
| sd(logAsym_Intercept) | 0.42 | 0.03 | 0.37 | 0.48 | 1.01 | 414 | 986 |
| sd(logtmid_Intercept) | 0.33 | 0.02 | 0.29 | 0.38 | 1.00 | 442 | 742 |
| sd(logslope_Intercept) | 0.16 | 0.01 | 0.14 | 0.18 | 1.00 | 648 | 1216 |
| ~geno (Number of levels: 48) |  |  |  |  |  |  |  |
| sd(logAsym_Intercept) | 0.07 | 0.01 | 0.05 | 0.08 | 1.00 | 1390 | 2427 |
| sd(logtmid_Intercept) | 0.05 | 0.01 | 0.04 | 0.06 | 1.00 | 1365 | 2506 |
| sd(logslope_Intercept) | 0.04 | 0.00 | 0.03 | 0.05 | 1.00 | 2029 | 3698 |
| ~geno:Env (Number of levels: 5952) |  |  |  |  |  |  |  |
| sd(logAsym_Intercept) | 0.05 | 0.00 | 0.05 | 0.05 | 1.00 | 2860 | 5605 |
| sd(logtmid_Intercept) | 0.01 | 0.00 | 0.01 | 0.01 | 1.00 | 1764 | 3395 |
| sd(logslope_Intercept) | 0.00 | 0.00 | 0.00 | 0.00 | 1.00 | 2708 | 3828 |
| Population-Level Effects: |  |  |  |  |  |  |  |
| logAsym_Intercept | 2.21 | 0.04 | 2.14 | 2.28 | 1.01 | 186 | 419 |
| logtmid_Intercept | 2.09 | 0.03 | 2.02 | 2.15 | 1.03 | 119 | 182 |
| logslope_Intercept | 1.29 | 0.01 | 1.26 | 1.32 | 1.01 | 303 | 653 |
| Family Specific Parameters: |  |  |  |  |  |  |  |
| sigma | 0.18 | 0.00 | 0.17 | 0.18 | 1.00 | 5237 | 4733 |

Draws were sampled using sample(hmc). For each parameter, Bulk\_ESS and Tail\_ESS are effective sample size measures, and Rhat is the potential scale reduction factor on split chains (at convergence, Rhat = 1).

We computed correlations between each of the DE parameter 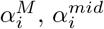 and 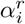 (estimated by posterior medians) on the one hand, and the individual APSIM parameters, on the other. We also obtained the asymptotic 95% confidence intervals based on the Fisher z-transformation using the standard **R** command **cor.test**. The results are presented graphically in Figure 10.

**Figure 10:**
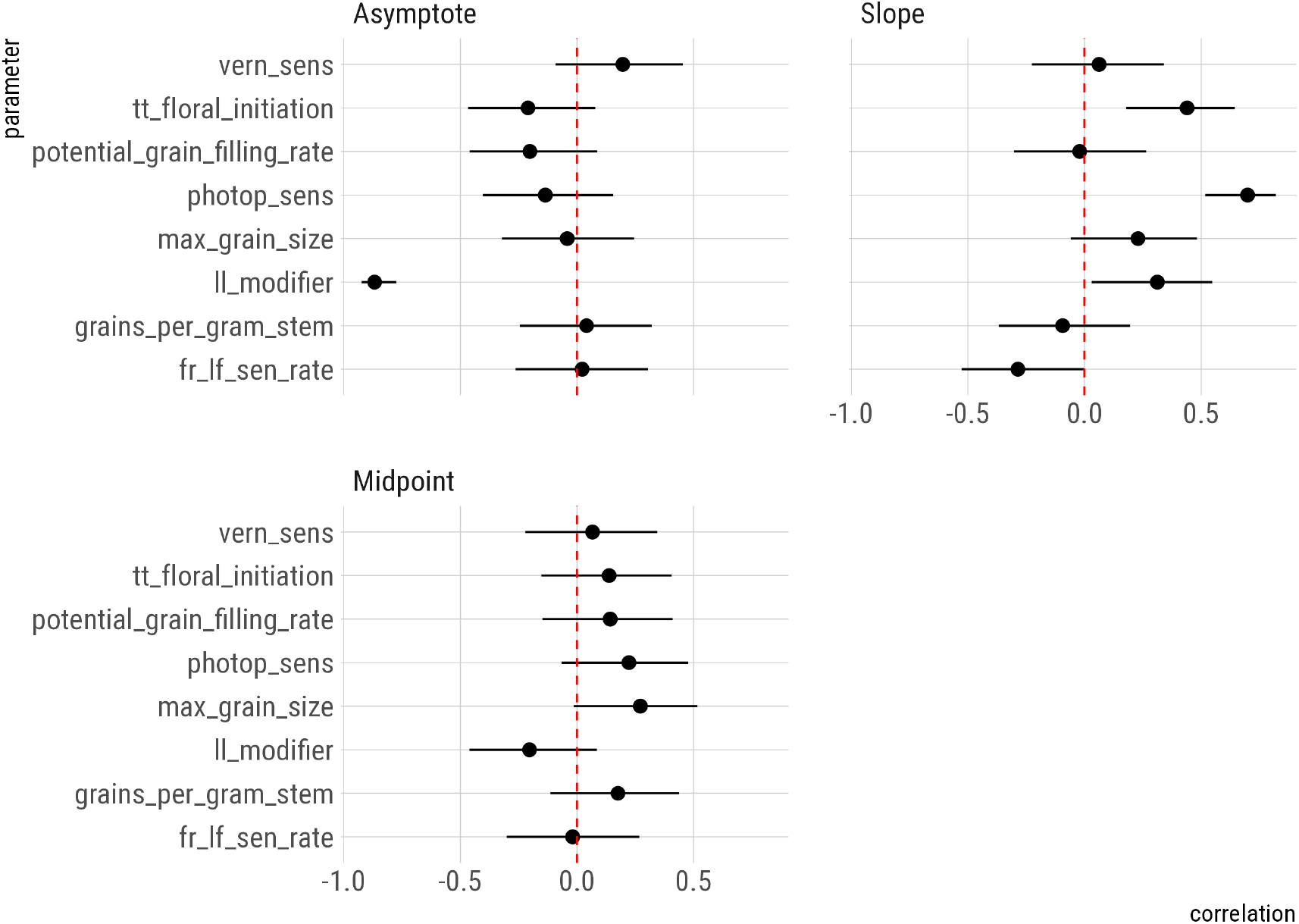
Correlations between APSIM parameters, on the one hand, and the genotype main effects 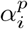, on the other, for superscripts *p* = *M*, *mid* and *r*. APSIM parameter labels are explained in Table 4. Each panel gathers results for a different parameter of the log-logistic function, identified by labels `Asymptote`, `Midpoint` and `Slope`; see Section 2.2.2 for the meaning of the parameters. Points correspond to estimates, while segments are 95% asymptotic confidence intervals based on Fisher’s z-transformation. The dashed red line has been added for reference to indicate zero correlation.

Several APSIM parameters show significant correlations with the inferred parameters of our Bayesian hierarchical model. The maximum biomass seems to be strongly negatively related to the lower limit for water uptake (`ll _modifier`), whereas that relationship is positive for the slope parameter. Higher slopes were associated with larger thermal times till floral initiation (`tt_floral_initiation`) and stronger photoperiod sensitivity (`photop_sens`). The midpoint time did not show any obvious correlation with APSIM parameters.

## 4 Discussion

### 4.1 Incorporating phenotyping information into mixed models for genomic prediction

In recent years, automatic monitoring of longitudinal phenotyping data has become an integral part of many plant breeding programs and experiments. Incorporating this kind of information into breeding strategies and decisions is a topic of current research. From a prediction perspective, genomic prediction models can be improved by combining genomic and phenotyping data into genotype-to-phenotype models. An important objective of such integrated models is to increase the predictive ability of within-trial phenotypes at later time points from genomic information and phenotyping information at earlier time points. In other words, the target phenotypes, such as yield, phenological traits, and yield components, are forecasted from early season information. The chosen modeling framework is often that of multi-trait mixed models and random regressions (Baba et al., 2020; Campbell et al., 2018; Momen et al., 2019; Sun et al., 2017, 2019). These models can contain complex formulations for the variance-covariance structures dealing with the correlations between traits over time and the incorporation of one or more kernels on the genotypes to represent genomic relations, but they do not include explicit model terms or structures that represent physiological mechanisms or insights. Furthermore, the above papers leave open how genomic and phenotyping information should be combined to predict yield in future environments under *G* × *E*.

Jarquín et al. (2014) presented a mixed model framework for genomic prediction between trials in the presence of *G* × *E* based on the definition of kernels (relationship matrices) for genotypes and environments. A generalization of this approach addresses *G* × *E* while integrating genomic and phenotyping information on genotypes with enviromic information on environments (Adak et al., 2023; Robert et al., 2022). Although the emphasis for this new type of integrated mixed models is on between-trial prediction in place of within-trial prediction, the representation of the additional phenotyping information inside the prediction models ignores most of the physiological principles that make yield the outcome of growth and developmental processes. In this approach, the dynamic environmental inputs that are typical for physiological models are not used. The totality of the physiological information is reduced to a relationship matrix between genotypes.

### 4.2 Crop growth model – whole genome prediction

In stark contrast to the mixed models above that incorporated phenotyping information without reference to physiological models, an alternative approach, called crop growth model - whole genome prediction, builds on elaborate physiological models for prediction in the expectation that such an approach will lead to better between-trial predictions of yield under *G* × *E* (Technow et al., 2015; Messina et al., 2018). The CGM-WGP approach predicts the end-point trait yield together with dynamical yield components for new genotypes and environments using marker profiles and dynamic environmental data as inputs. The marker profiles serve to provide a molecular basis to a limited number of physiological traits. Marker effects are estimated following a Bayesian procedure. The CGM-WGP represents a powerful, general and attractive predictive framework founded on prior physiological knowledge. In its initial formulations, the

CGM-WGP approach does not use dynamic observations on plant growth during the season and has to rely fully on the correctness of the pre-specified parameters and relations inside the crop growth model that converts marker profiles and environmental inputs into yield outputs.

Crop growth models tend to have many parameters and it is impossible to estimate all of the parameters in crop growth models at a genotype-specific level. When using a CGM-WGP approach, a decision has to be made about which parameters to allow to vary with genotype and which to fix for the crop species as a whole. In the CGM-WGP papers by Technow et al. (2015), Cooper et al. (2016), Messina et al. (2018) and Jighly et al. (2023) only a few yield components are genotype specific and a function of marker inputs. Although the CGM-WGP approach is dynamic for the inputs from the environment and the outputs of the yield components, it does not use dynamic inputs from phenotyping and phenomic devices. Such inputs may contain important information on the occurrence of biotic and abiotic stresses.

### 4.3 Our hierarchical dynamic model proposal to function-valued modeling of longitudinal traits in multiple trials

The Bayesian framework we developed in this paper provides a flexible framework to model a longitudinal trait for a set of genotypes across multiple environments, taking into account *G* × *E*. Our approach is unique because it fits the growth curves simultaneously for all included genotypes and environments. Therefore, propagation of errors and tests about differences between growth trajectories can be performed within a well-defined context, namely that of the Bayesian framework that relies for inference on the properties of the posterior distribution. The same framework contains the tools for constructing prediction intervals along the fitted growth trajectories. Each trajectory is determined by a limited number of biologically interpretable DE parameters whose expectation is itself a GLMM with main effects for genotype and environment, as well as *G* × *E*.

In our approach, we followed a function-valued trait modeling perspective to growth trajectories as propagated by Wang et al. (2014), Gomulkiewicz et al. (2018) and Baker and Wang (2021). Within that framework, we interpret the DE parameters as phenotypic traits themselves, as traits underlying the dynamics of the yield component or secondary trait that we are studying. Analogously to a quantification of the sources of variation in the context of linear mixed models in quantitative genetics, the contributions of genotype, environment, and *G* × *E* to the total ‘phenotypic variation’ for the DE parameter can be assessed within the Bayesian framework by inspecting the posterior distributions for the corresponding dispersion parameters. Their magnitudes reflect their relative importance. For classical phenotypes such as yield, biomass and phenology, the environment is by far the most important source of variation, usually followed by the genotype when the environmental conditions are not too wildly different between trials and finally the *G* × *E*. Exactly that order of importance also occurred for the DE parameters in our model for biomass, for all three the DE parameters of asymptote, midpoint, and slope. Comparison of dispersion parameters between the three DE parameters identified the asymptote as the parameter with the strongest *G* × *E*, that is, with the highest ratio of *G* × *E* to genotype variation. For the midpoint and slope, *G* × *E* was small to absent, which means that the dynamics of biomass followed an additive model for these two parameters.

### 4.4 Alternative function-valued modeling approaches for longitudinal traits

A function-valued approach to plant growth for populations of genotypes was also proposed by Baker et al. (2018a,b). For leaf width and length in *Brassica rapa*, logistic models were fitted on growing degree days per plant. The growth parameters were then extracted and subsequently further analyzed in a Bayesian hierarchical analysis of variance (ANOVA) model containing a population, genotypic and plant level. Both Baker et al. (2018a) and Baker et al. (2018b) tested plant growth under two plant densities that were interpreted as different environmental conditions. A test for genotype by density interactions was performed and on failure to reject the null hypothesis of no interaction, it was concluded that the growth of a genotype as modeled under one density could be used to predict the growth of the same genotype under an alternative density. In our approach, in contrast to Baker et al. (2018a) and Baker et al. (2018b), a single inferential framework was used for both modeling the dynamics of plant growth and expressing the dynamic parameters in terms of genotype, environment, and *G* × *E*. A test of *G* × *E* for individual DE parameters followed straightforwardly from inspection of the posterior for the corresponding dispersion parameter.

Baker et al. (2018a) extended their Bayesian ANOVA model for the logistic parameters by making the genotypic means for these parameters dependent on photosynthetic capacity. In this way, they wanted to test whether carbon limitations were responsible for genotypic differences as a factor affecting the growth dynamics of individual genotypes. The photosynthetic capacity can be measured in different ways, some of them being very time consuming others delivering approximations via remote sensing. The insertion of approximate physiological parameters from phenotyping devices or phenomic measurements to describe variation in growth parameters offers an attractive and powerful modeling perspective. Similar tests for DE parameters being a function of certain genotype-dependent limitations or capacities can naturally be implemented in our framework by introducing genotypic covariates into the GLMMs for the DE parameters and inspecting the posterior distribution for the slope with respect to such a covariable.

Baker et al. (2018b) identified quantitative trait loci (QTLs) for dynamic parameters, after estimation of those dynamic parameters in logistic growth models. They follow up by creating QTL-based predictions for the full growth trajectories. Again, this is a multi-step approach. Within our Bayesian hierarchical model for plant growth, we can in principle directly predict the dynamical response from marker profiles by replacing the genotypic main effects in the GLMMs for the growth parameters by penalized regressions on marker profiles. With a double exponential prior on the marker effects this would lead to QTL mapping via LASSO (Least Absolute Shrinkage and Selection Operator) regression (Li and Sillanpää, 2012), while with a normal prior we would arrive at genomic prediction using a ridge regression.

In the field of animal breeding, Varona et al. (1999) use the Gibbs sampler (or, rather, the Metropolis-within-Gibbs sampler) with an explicit form of the growth curve, specifically the Von Bertalanffy growth curve, within the mixed effects modelling framework. Their approach is not generalisable to the differential equation models that do not admit explicit solution. The Gibbs sampler also does not scale well in such complex nonlinear models. In fact, Onogi et al. (2019), who use a similar computational approach as Varona et al. (1999), are constrained to run their Gibbs sampler for millions of iterations (Onogi et al., 2019, p. 5). This is indicative of slow mixing of the sampler. In contrast, in order to efficiently explore the posterior distribution, a few thousand iterations typically suffice with the NUTS sampler (Hoffman and Gelman, 2014; Betancourt, 2017; Betancourt et al., 2017), which is the computational algorithm of **brms**. The mathematical model in Onogi (2020) is similar to that in Varona et al. (1999); Onogi et al. (2019) and does not envision a governing differential equation without a closed-form solution. The computational approach they use is a hybrid method from Onogi et al. (2015). This combines Gibbs sampling with coordinate ascent variational inference (CAVI). However, even though in general faster than MCMC sampling, CAVI is closely related to Gibbs sampling and suffers from the same general drawbacks (Blei et al., 2017). A computational bottleneck of the Bayesian approach to inference in DE models lies in the need for their repeated numerical integration in likelihood evaluations. Already for a medium complexity model, this is not realistically feasible when the computational method requires millions of iterations until convergence (Voit and Almeida, 2004, Appendix 1). Note also that while the MCMC methods are guaranteed (under idealised conditions) to recover the posterior distribution as the number of iterations grows indefinitely (Roberts and Rosenthal, 2004), this is not the case with variational inference, which involves a non-vanishing approximation error even in the limit.

### 4.5 Outlook

The current version of our hierarchical Bayesian model for longitudinal traits in multiple trials is promising as a starting point for a framework for the prediction of yield under *G* × *E*. However, a number of issues require attention. First of all, from the candidate DE models described in Van Voorn et al. (2023) we chose for this paper the simplest one, the logistic model, which was slightly modified, for computational reasons, into a log-logistic model. The log-logistic model does not contain an environmental driver, as in the irradiance model or the temperature model of Van Voorn et al. (2023). Neither does the log-logistic model contain an environmental state variable, as in the water model of Van Voorn et al. (2023). The log-logistic model has an explicit solution that turns our hierarchical dynamic model into a hierarchical regression model. Therefore, estimation procedures are obviously easier to perform. Nevertheless, the strong point of our approach remains that parameters for growth processes can explicitly be modeled as functions of genotypic, environment and *G* × *E* terms. The usefulness of parameters of yield components for incorporation into predictive models for yield will firstly depend on the extent to which these parameters are free of *G* × *E*. Even when parameters for yield components do show *G × E*, they can still be useful for yield prediction as long as the *G* × *E* can be modeled in terms of environmental covariables, that is, the *G* × *E* term for the rate parameter is written as the product of a genotypic sensitivity times a predictable environmental condition. The situation here is comparable to the use of factorial regression in models for *G* × *E* in target traits (Van Eeuwijk et al., 2016; Avagyan et al., 2025).

When our DE system does not have an explicit analytical solution, our hierarchical Bayesian framework will have to solve the DE system for each combination of genotype and environment numerically together with estimating the GLMM parameters for the DE parameters. This is possible within our modeling framework that relies on the Bayesian modeling language **Stan** and we will present results in follow-up papers. Computing resources will need to be increased when there is no explicit solution for the DE system. For the analyses presented in this paper, we kept the playing field small as the objective was to investigate the principles of a hierarchical Bayesian approach to longitudinal modeling. A first next step will consist of upscaling our modeling facilities to medium sized panels of genotypes (100s) and sets of environments (25–100).

An ultimate aim of longitudinal traits modeling in phenotyping experiments is to arrive at a predictive model that allows us to predict yield under *G* × *E*. Different DE systems can be tried to see whether formulations can be found in which parameters have no or limited *G* × *E* or in which *G* × *E* depends on predictable environmental covariables. Further, the genotypic terms, intercepts, and slopes in the GLMMs for the DE parameters should be predictable from marker profiles, like in standard genomic prediction approaches. For the prediction of genotypic intercepts and slopes for DE parameters, an additional layer can be added to the Bayesian hierarchy. An alternative is a two-step approach in which these genotypic intercepts and slopes are predicted in a separate genomic prediction step. An interesting aid in identifying candidate DE systems for modeling longitudinal traits is the SINDy-PI algorithm (Kaheman et al., 2020) that tries to identify sparse representations of dynamic systems. An illustrative application of this approach in the context of genotype-specific crop growth modeling is presented by Verburg et al. (2026). We feel that our hierarchical Bayesian approach to longitudinal trait modeling opens up new perspectives to predict yield under *G* × *E*. This approach provides an additional tool for breeders and geneticists in conjunction with the mixed kernel-based models following the strategy outlined by Jarquín et al. (2014) and the CGM-WGP strategy introduced by Technow et al. (2015) and Messina et al. (2018).

## Author contributions

Modelling **G** × **E** × **M** was conceived between Corteva Agriscience and Wageningen University & Research – Biometris, with groups led by CM and FV, respectively. Conceptualization and development was done by all authors. The APSIM wheat simulation data were created by DB. Model fitting was done by SG. The first version manuscript writing and preparation was done by SG, GV and FV. Intermediate editing of the manuscript was done by all authors. SG, GV, MB and FV did the final manuscript editing. All authors contributed to the article and approved the submitted version.

## Funding

This research was supported by Predictive Agriculture, Research & Development at Corteva Agriscience (2019–2020).

FV worked on this paper partly in the context of the EU-INVITE project, which is part of the Horizon 2020 Framework Programme of the European Union, under Grant Agreement 817970 (https://www.h2020-invite.eu/).

FV, MB and SC contributed to this paper partly within the framework of the GRDC INVITA UQ-WUR and GRDC AAGI UQ-WUR Collaborative Research Agreements.

SG, MB and GV acknowledge funding from the WUR Investment Theme on Data-Driven Discoveries in a Changing Climate (D3-C2).

DB contributed to this paper thanks to the funding of Subvención a la Instalación en la Academia, ANID 85220037.

## Acknowledgments

Acknowledgment is made to the APSIM Initiative, which takes responsibility for quality assurance and a structured innovation program for APSIM’s modelling software, which is provided free for research and development use (see https://www.apsim.info/ for details).

## Data availability statement

The APSIM wheat dataset analyzed in this study can be accessed via Bustos-Korts (2023).

## Declaration of AI use

Parts of the **R** analysis script were written with the assistance of GitHub Copilot inline code completions in RStudio 2024.12 during 2025. The completions were generated by GitHub’s default completion models at the time: a GPT-3.5 Turbo-based model, GPT-4o Copilot and GPT-4.1 Copilot. The authors reviewed, tested and revised all code suggested by Copilot and verified that the results are accurate and reproducible. Generative AI was not used to design the model, interpret the results, or write the manuscript. AI assistance (Claude Opus 5.5) was utilized during the final preparation of the submission strictly for language enhancement, stylistic refinement, and copy editing.

## Software

The computer code used in this paper is available through the dedicated GitHub repository at https://github.com/gugushvili/dynamic_phenotypes.

## Appendix A Supplementary Figures

**Figure 11:**
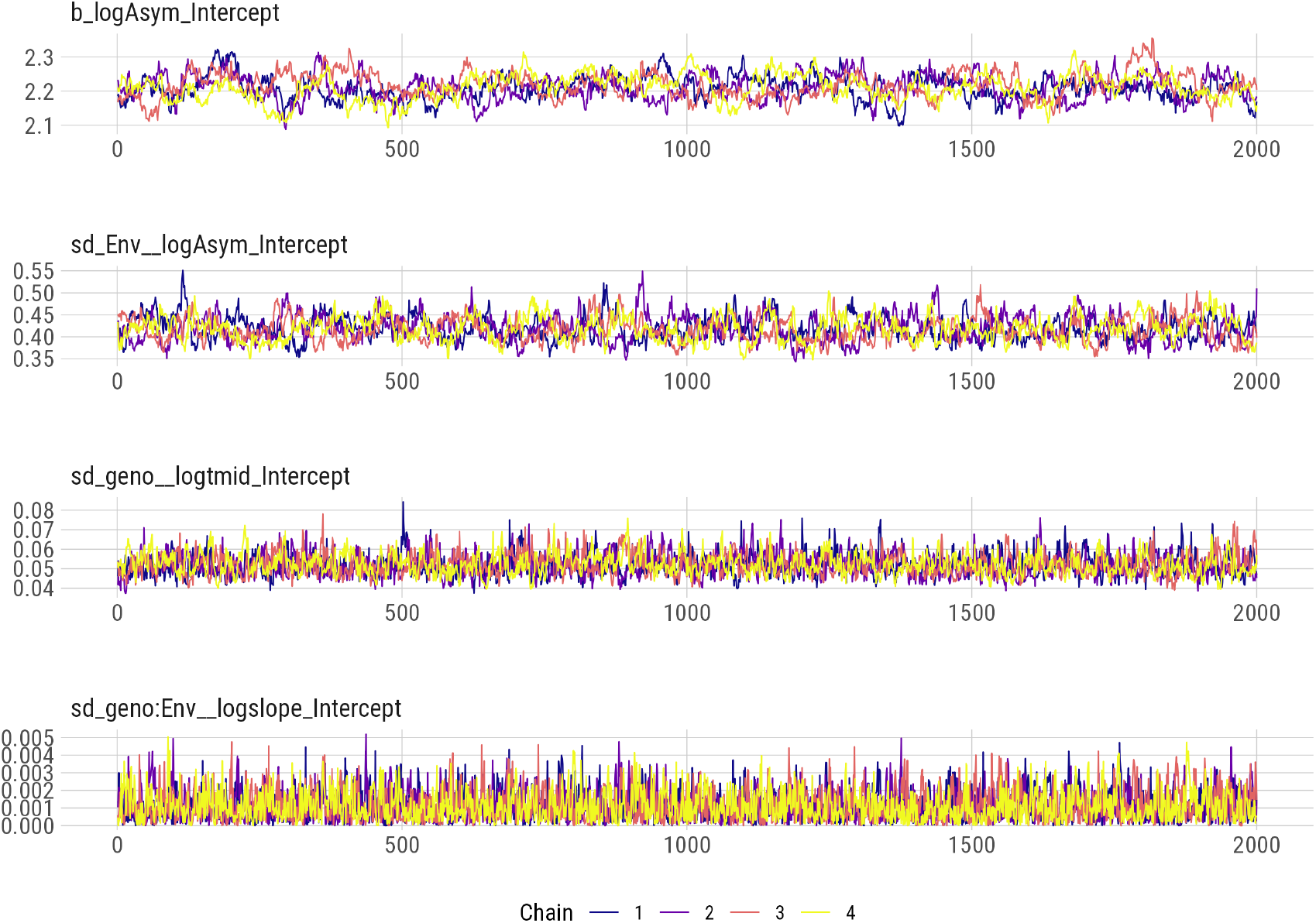
Trace plots for several model parameters. There are 4 Markov chains per parameter.

**Figure 12:**
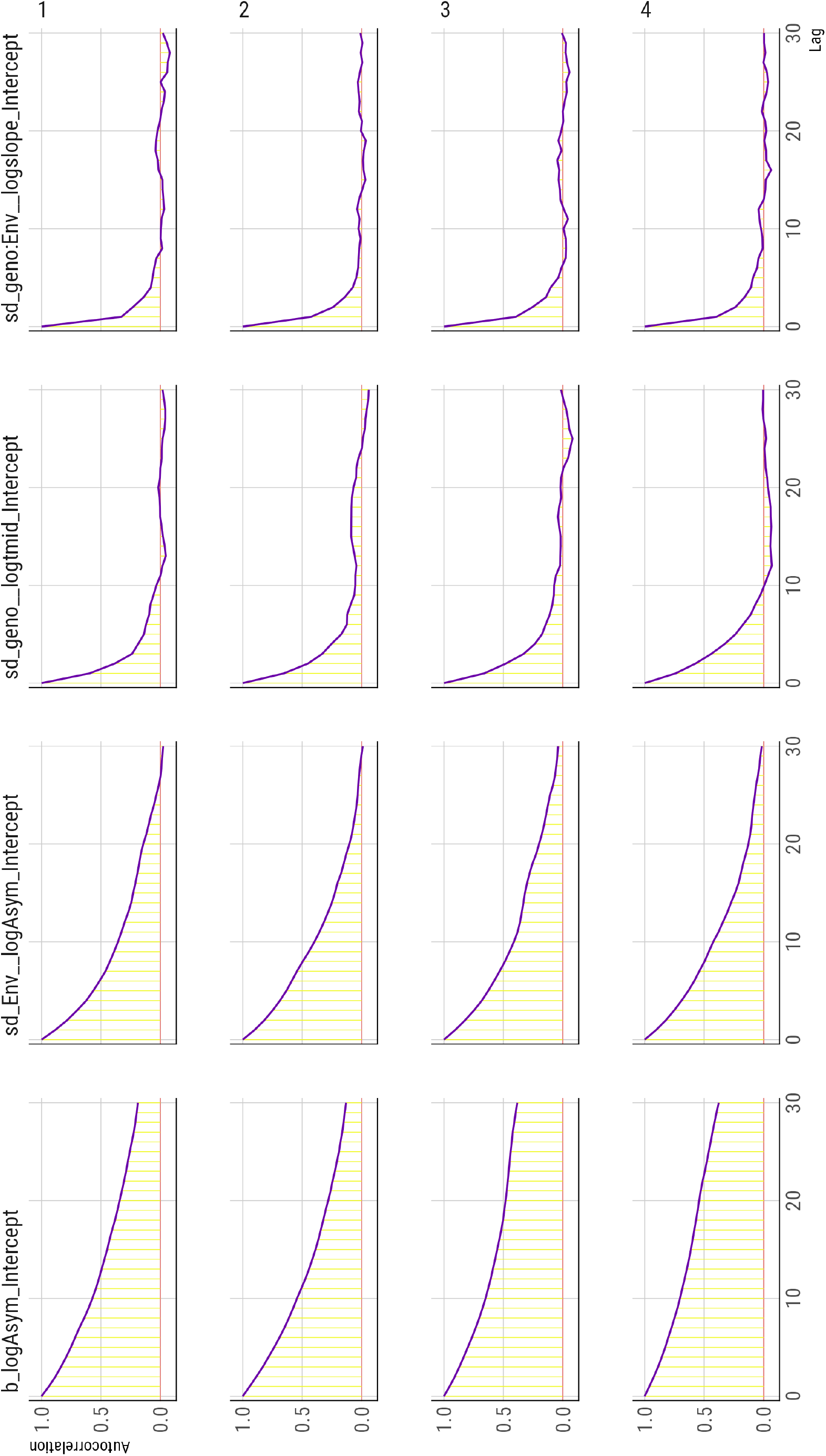
The autocorrelation function for Markov chains of several model parameters. The figure is best viewed sideways. There are 4 Markov chains per parameter. Each is identified by a number from 1 to 4.

**Figure 13:**
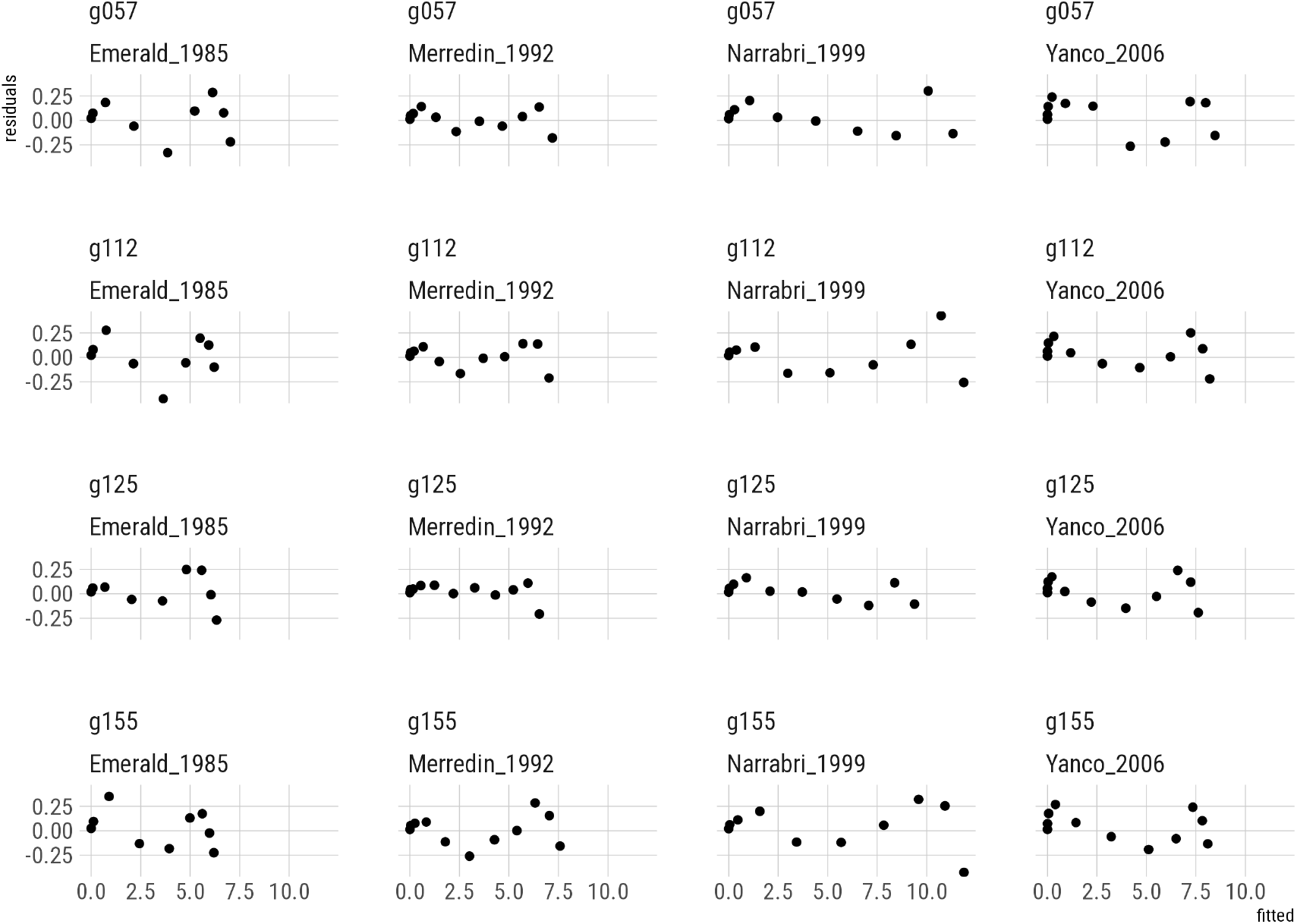
The residuals versus fitted values plots for an arbitrary selection of 4 genotypes in 4 trials.

## Appendix B Supplementary Listings and Tables

**Listing 1:**
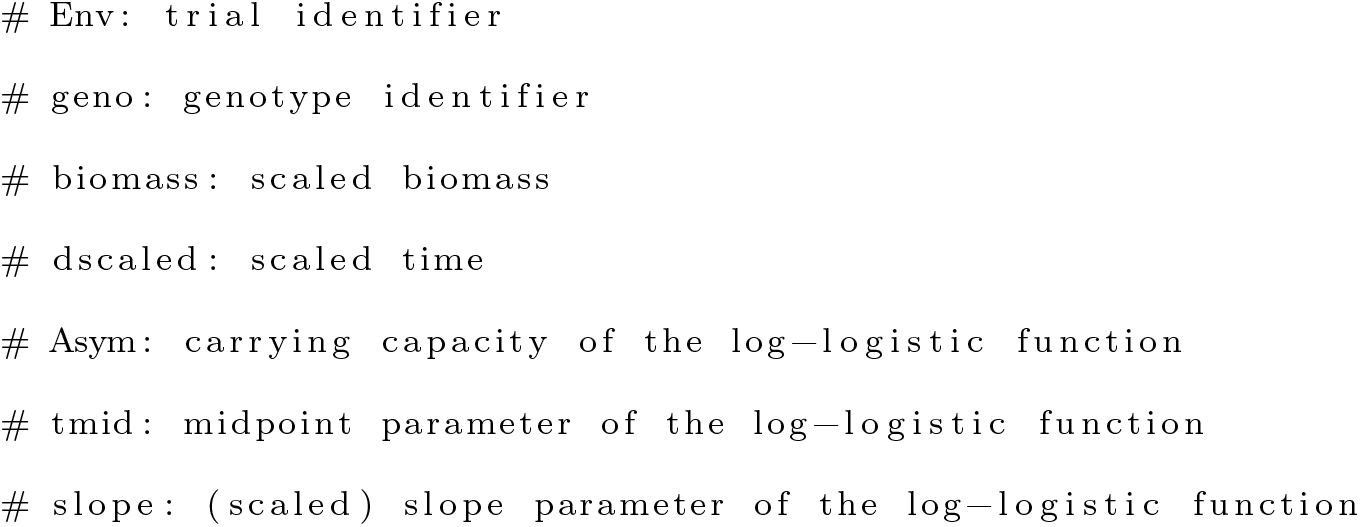
Variable and parameter names in the **brms** model.

**Listing 2:**
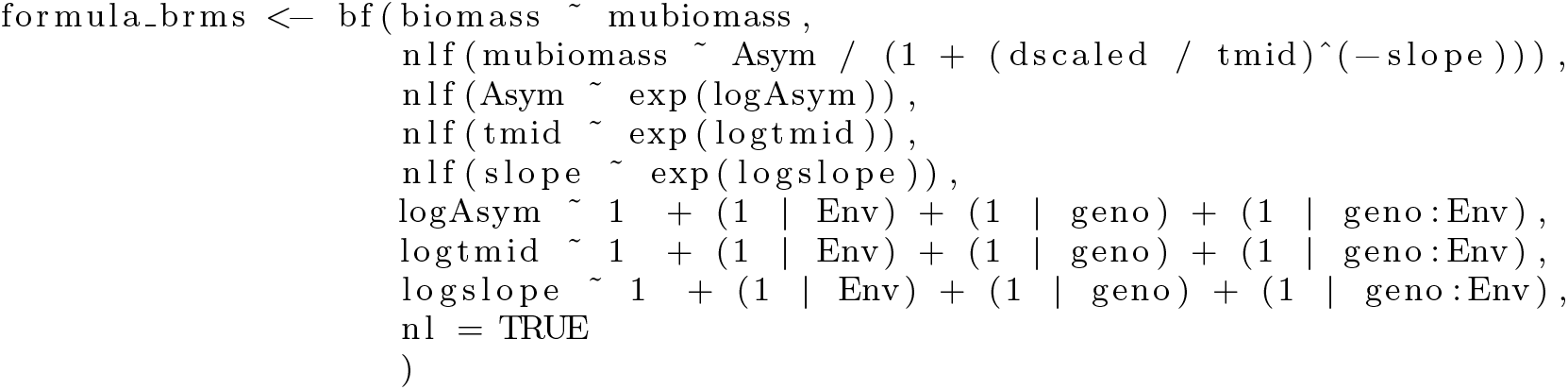
Model formula as implemented in **brms**.

**Listing 3:**
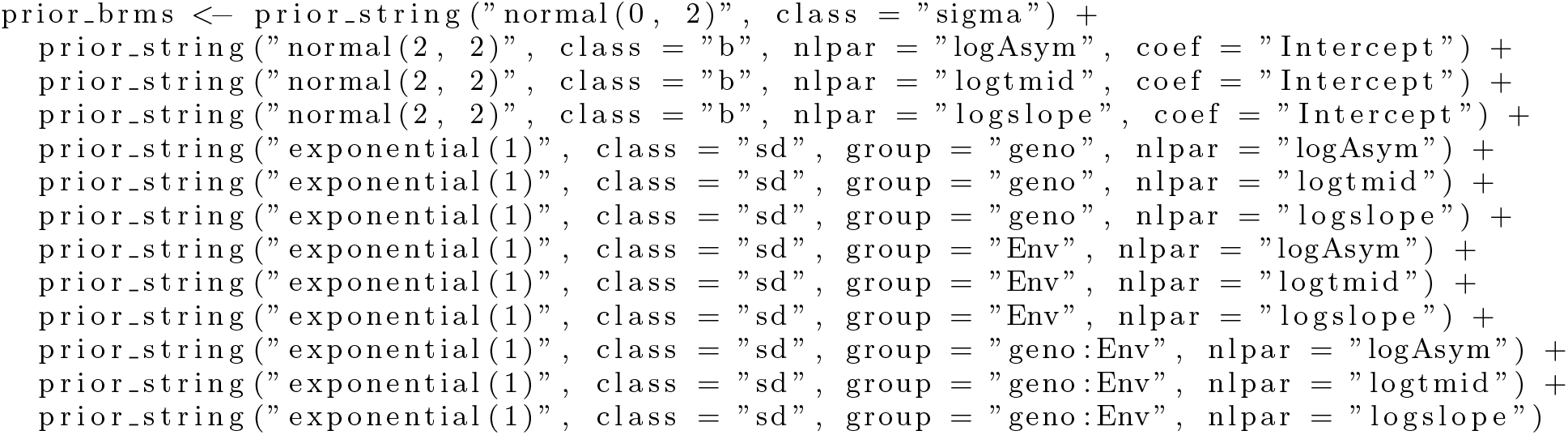
Priors for the **brms** model.

